# The complement system supports normal postnatal development and gonadal function in both sexes

**DOI:** 10.1101/233825

**Authors:** Arthur S. Lee, Jannette Rusch, Abul Usmani, Ana C. Lima, Wendy S.W. Wong, Ni Huang, Maarja Lepamets, Katinka A. Vigh-Conrad, Ronald E. Worthington, Reedik Mägi, John E. Niederhuber, Xiaobo Wu, John P. Atkinson, Rex A. Hess, Donald F. Conrad

## Abstract

Male and female infertility are clinically managed and classified as distinct diseases, and relatively little is known about mechanisms of gonadal function common to both sexes. We used genome-wide genetic analysis on 74,896 women and men to find rare genetic variants that modulate gonadal function in both sexes. This uncovered an association with variants disrupting *CSMD1*, a complement regulatory protein located on 8p23, in a genomic region with an exceptional evolution. We found that *Csmd1* knockout mice display a diverse array of gonadal defects in both sexes, and in females, impaired mammary gland development that leads to increased offspring mortality. The complement pathway is significantly disrupted in *Csmd1* mice, and further disruption of the complement pathway from joint inactivation of *C3* leads to more extreme reproductive defects. Our results can explain a novel human genetic association with infertility and implicate the complement system in the normal development of postnatal tissues.

## Introduction

Male and female infertility have historically been classified and clinically treated as distinct disease entities and this perspective has led to the assembly of many cohorts for the study of sex-specific reproductive processes (Hotaling and Carrell, 2014; Nelson, 2009; O’Flynn O’Brien et al., 2010; Stolk et al., 2012). However, many molecular and physiological mechanisms of fertility regulation are shared between male and female mammals including embryonic gonad development, meiosis, and the hypothalamic-pituitary-gonadal axis (Matzuk and Lamb, 2008). There are other phenomena common to gonadal function in both sexes that are poorly understood, and the extent to which these phenomena have a common set of regulators is unknown. For instance, programmed germ cell degeneration is a pervasive part of gonadal biology in both sexes. In human males, roughly 80% of the meiotic descendants of spermatogonial stem cells undergo apoptosis prior to ever becoming spermatozoa (Hess and Renato de Franca, 2008). In human females, nearly 80% of the oocytes made during embryogenesis are eliminated by birth, representing the first major stage of oocyte loss (Baker, 1963; Kurilo, 1981). Upon menarche, a woman will ovulate approximately 400 times in her life.

However, of 300,000-500,000 oocytes present at birth, only roughly 1,000 survive the sojourn to menopause, representing colossal germ cell loss not attributable to ovulation (Wallace and Kelsey, 2010). The mean ratio of surviving: apoptotic germ cells differs between species, but is narrowly regulated within species (Hsueh et al., 1994) (Hess and Renato de Franca, 2008). Germ cell loss in both sexes may represent a cellular safeguard against violation of essential cellular events such as DNA replication/repair and chromosome segregation--events that occur prior to, or during, meiosis. Spermiogenesis and folliculogenesis, which occur after the onset of meiosis, are highly complex in their own right. Molecular mechanisms for error-checking these processes are poorly understood.

Defects in the development of germ cells that are due to problems originating in the gonad are clinically defined as primary gonadal dysfunction. Primary gonadal dysfunction is an infertility phenotype that is attractive for human genetic analysis, has a prevalence of at least 1% in males and females (Luborsky et al., 2003; Willott, 1982), and has clear diagnostic criteria. In males, primary gonadal dysfunction can manifest as a total absence of germ cells, an arrest of spermatogenesis, or complete but limited sperm production. In females the presentation can range from complete absence of germ cells to irregular ovulation or premature menopause.

We have previously identified a reproducible association between rare copy number variant (CNV) burden and male gonadal dysfunction (Huang et al., 2015; Lopes et al., 2013). In the present study, we used array and exome sequencing data from a large cohort of post-menopausal women, collected as part of the Women’s Health Initiative study (Chen et al., 2012), to identify novel, shared factors required for normal gonadal function in both sexes, and replicated our findings with data from the UK Biobank(Sudlow et al., 2015).

## RESULTS

### Rare *CSMD1* mutations are associated with reproductive outcomes in humans

Due to the strong selective pressure against infertility mutations, we hypothesized that male and female gonadal dysfunction are driven largely by rare mutation events. To test this hypothesis, we acquired SNP array and phenotype data from 12,002 women (515 cases of inferred primary ovarian insufficiency (POI) vs. 11,487 normal menopause controls) and 2,072 men (321 cases of spermatogenic impairment vs. 1,751 controls) with known reproductive health history. Since it is difficult to detect rare variants via conventional SNP arrays, we leveraged the SNP log R ratios and B-allele frequencies to discover CNVs that occupy the entire allele frequency spectrum (**Table S1, Methods**). We then applied filters to enrich for deleterious CNVs (minor allele frequency < 0.01 and length > 100 kb). We used these CNVs to perform a rare variant, gene-based, case-control genome wide association study (GWAS) separately in males and females (**Methods**).

Our rare variant GWAS identified a significant association between inferred POI and deletions overlapping the CUB and Sushi multiple domains 1 (*CSMD1*) gene located on chromosome 8p23.2 (OR = 16; nominal p-value=4.0 × 10^−4^; genome-wide p-value= 0.015; **Figure 1A**). This association signal replicated in our smaller cohort of male spermatogenic impairment (OR = 3.3; nominal p-value = 6.5 × 10^−3^). This CNV association is largely driven by the observation of an aggregate enrichment of rare deletions in cases, compared to controls, all of which are clustered in the 5’ half of the gene, in introns 1-3 (**Figures 1B**) There was no single CNV in the region with a significant frequency difference between cases and controls.

**Figure 1.**
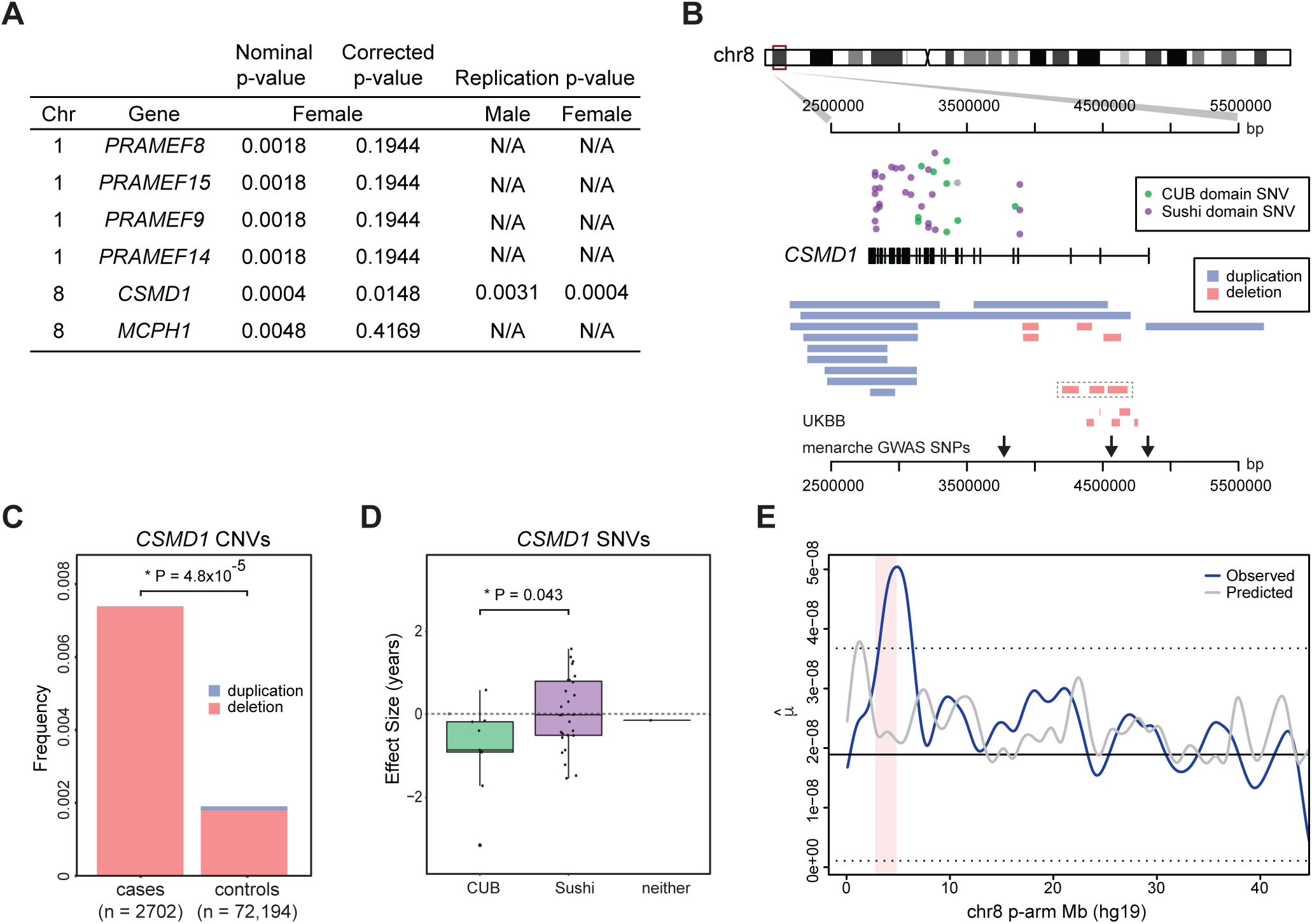
The landscape of rare mutations across *CSMD1* in humans. **(A)** Rare CNVs associated with gonadal function. We performed a gene-based genomewide association study to identify rare CNVs associated with female gonadal dysfunction (stage I) and attempted replication in males (stage II). All genes with nominal associations p-values < 0.05 in stage I, and the analogous values for stage II are listed; a value of “N/A” indicates that no CNVs were observed at the locus. Of note, biallelic knockout of *Mcph1* was reported to cause infertility in male and female mice (White et al., 2013). **(B)** Location of 37 rare (MAF < 0.01) SNVs overlapping *CSMD1* among 1,526 exome-sequenced females, and large, rare CNV regions overlapping *CSMD1* among 14,074 females and males (836 cases and 13,238 controls). CNV regions found in males are outlined by a dashed box. Additional tracks: “UKBB”, the location of all rare intron 1-3 deletions observed in the UK Biobank POI cohort – for clarity only the deletions observed in cases are shown; “menarche GWAS SNPs”, the location of three statistically independent lead SNPs from a large-scale GWAS of age at menarche are depicted as arrows along the bottom of the figure: rs2688326, rs 2724961, and rs4875424 (Day et al., 2017). **(C)** Stacked barplot depicting frequency of rare (MAF < 1%) CNVs overlapping introns 1-3 of *CSMD1* among 2,702 cases of male or female gonadal dysfunction versus 72,194 controls. Rare deletions over *CSMD1* segregate significantly with cases (OR = 4.09; meta-analysis p = 4.8 × 10^−5^). **(D)** Boxplot depicting the effect size of rare *CSMD1* SNVs found in females, stratified by protein domain (CUB, Sushi, or neither). SNVs occurring in the CUB domains are significantly associated with an earlier onset of menopause when compared to SNVs in the Sushi domains (β_CUB_ = -0.86, 95% CI [-1.56, -0.151]; β_SUSHI_ = 0.046, 95% CI [-0.255, 0.377]; P = 0.043; Wilcoxon rank-sum test). **(E)** *De novo* mutation (DNM) frequency across chromosome 8. DNMs were called from whole genome sequence data for 709 parent-offspring trios. DNMs were compiled across 100 kb windows across chromosome 8 and a smoothing spline function was applied to the data (blue line). We used a mutation-rate prediction model to estimate context-dependent mutation rates for all bases on chromosome 8, averaged these across fixed 100 kb windows (grey line). Solid horizontal lines represent the mean value across chromosome 8, dotted horizontal lines represent 1 standard deviation about the mean, and the pink shaded region represents the interval encompassing *CSMD1.μ̂*, the estimated germline mutation rate per base-pair, per generation.

To replicate the association between deletions in *CSMD1* and risk for gonadal dysfunction, we constructed another POI case-control cohort using the UK Biobank (**Methods**). After CNV QC and rigorous case/control selection, we obtained a cohort of 63,064 women with both reliable phenotype data and CNV calls; 1,873 of these were considered cases of POI. We again observed a significant association between POI and rare deletions in introns 1-3 of *CSMD1* (0.6% frequency in cases, 0.2% in controls, OR=3.03, p< 5 × 10^−4^, Figure 1A and 1B). To succinctly summarize the risk conferred by rare (<1% MAF) deletions in introns 1-3 of *CSMD1,* we performed a meta-analysis across all three cohorts, this time considering deletions of all sizes, and found a frequency of 0.7% in cases and 0.2% in controls (meta-analysis p = 4.8 × 10^−5^; **Figure 1C**). The list of deletions observed in introns 1-3 are provided as **Table S2**.

To further replicate our findings using an orthogonal genotyping platform, we analyzed single nucleotide variants (SNVs) ascertained by whole-exome sequencing generated from the female cohort (n = 1,526). Employing SKAT, a gene-based quantitative trait association framework, we identified a significant association between rare (MAF < 0.01), deleterious *CSMD1* single nucleotide variants and age at menopause (p-value < 5 × 10^−3^; **Methods**). The bulk (97.1%) of the *CSMD1* protein product consists of alternating CUB (complement C1r/C1s, **<u>U</u>**egf, **<u>B</u>**mp1) and Sushi/CCP (**<u>c</u>**omplement **<u>c</u>**ontrol **<u>p</u>**rotein) domains. We used linear models to further partition the association signal among these two domains. The *CSMD1* SNV association was driven almost exclusively by rare, deleterious mutations in the CUB (β_CUB_ = -0.86), but not Sushi (β_SUSHI_ = 0.046) domains (P = 0.043; for difference in effect size; **Figure 1D**). We estimate that each rare, deleterious mutation that we detected in CUB domains of *CSMD1* accelerates the onset of menopause by 10 months. These results immediately cast light on the relative importance of the CUB domain in the etiology of infertility, and prioritizes a potential target domain for therapy. Finally, while this work was in progress, a well-powered common variant GWAS in a female cohort of 182,416 individuals identified 3 common SNPs over *CSMD1* to be significantly and independently associated with age at menarche: rs2688325 (p=2.1 × 10^−9^), rs7828501 (p=1.2 × 10^−13^), and rs7463166 (p=1.3 × 10^−8^) (Perry et al., 2014). These 3 associations were replicated in ∼300,000 individuals: rs2688326 (p = 4.34 × 10^−18^), rs2724961 (p = 3.76 × 10^−33^), and rs4875424 (p = 1.99 × 10^−16^) (Day et al., 2017). Remarkably, these 3 common variant associations co-localize to the same 1Mb window as the rare disease-associated deletions described above (**Figure 1B**). Subsequent work has shown that age at menarche and menopause are positively correlated and that the common variants in *CSMD1* associated with age at menarche correctly predicted age at menopause in the expected direction (β_rs2688325_ = 0.014 +/-0.023; β_rs7828501_ = 0.021 +/-0.020; β_rs7463166_ = 0.031 +/-0.021) (Day et al., 2015). In summary, we detected associations between rare variants in *CSMD1* and gonadal dysfunction i) across multiple classes of genetic variation; ii) ascertained by orthogonal genotyping platforms; iii) occupying multiple points along the allele frequency spectrum; and iv) in multiple populations and cohorts.

### The *de novo* mutation rate across *CSMD1* is exceptionally high in humans

Excluding the Y chromosome, the distal arm of chromosome 8p contains the region of the genome with the greatest intra-population nucleotide diversity and the greatest nucleotide divergence between human and chimpanzee (Nusbaum et al., 2006). This signal of diversity and divergence peaks over *CSMD1* in a 1 Mb region that was originally reported to have an average human-chimpanzee divergence of 0.032 substitutions/bp, or 8.6 s.d. above the genomic mean. Multiple, non-exclusive factors can influence nucleotide diversity at a locus, namely mutation rate, demographic history, and natural selection. To evaluate the effect (if any) of mutation rate separate from confounding factors such as demography and long-term selection, we measured directly the number of *de novo* mutations (DNMs) across chromosome 8 in 709 human parent-offspring trios, calculating the average mutation rate in non-overlapping 100kb windows (**Methods**). We observed a local enrichment of DNMs overlapping *CSMD1,* as the mutation rate in six of the twenty 100kb windows over the gene was estimated to be greater than 6 × 10^−8^ mutations/bp/generation, a five-fold increase above the genomic average of 1.2 × 10^−8^ (**Figure 1E**). The “hottest” mutation hotspot we observed in the region had a DNM rate of 1.48 × 10^−7^, at 3.9 Mb-4.0 Mb, located within the nexus of infertility risk mutations reported above. This enrichment of DNMs is not well-explained by the intrinsic mutability of the primary nucleotide sequence in this region (**Figure 1E; Methods**). Using an association study on the same cohort of trios, we tested the region for cis-acting variants that might predispose to genome instability and, as an indirect result, infertility, but were unable to find a replicable association (data not shown).

### CSMD1 is expressed at the interface of germ cells and somatic cells in male and female gonads

*CSMD1* encodes for an extremely large (>3,000 amino acid) transmembrane protein with a large extracellular portion consisting of alternating CUB and Sushi complement-interacting domains (Kraus et al., 2006). The protein encoded by *CSMD1* is conserved between human and mouse, with 93% amino acid identity and 100% identity of the number and ordering of CUB and Sushi domains (**Figure 2A**). *CSMD1* and its mammalian orthologs are expressed in both male and female gonads, but little is known of its molecular function, particularly in the context of fertility. To elucidate CSMD1’s function, we first performed RNA-seq on whole mouse testes and ovaries. *Csmd1* is expressed in both tissues (**Figure 2B**), consistent with previous work (Soumillon et al., 2013; Steen et al., 2013). In testes, *Csmd1* is minimally expressed at 20 days and more robustly expressed at 40 days of age which coincides with the onset of sexual maturity. Mammalian testes demonstrate exceptional transcriptional complexity in comparison to other tissues, owing to the highly coordinated spatial and temporal synchronization required for proper spermatogenesis (Soumillon et al., 2013). Therefore, to capture a detailed transcriptional profile of *Csmd1,* we purified individual germ cell types using FACS (**Figure S1A**). Subsequent RNA-seq of purified germ cells reveals low levels of *Csmd1* expression during the diploid cell stages (i.e., spermatogonia and primary spermatocytes), and peak expression at the haploid stages (i.e., secondary spermatocytes and spermatids) (**Figure 2B; Figure S1B**). Finally, *in situ* antibody immunofluorescence on testis cross sections using a validated antibody (**Figure S2**) demonstrates that CSMD1 protein is expressed at the cell membrane at multiple stages of spermatogenesis, including at the interface of elongated spermatids and Sertoli cells, but is absent from spermatozoa, consistent with mRNA expression data (**Figure 2C; Figure S1C**). We performed immunofluorescence (IF) staining for key markers on whole-mount longitudinal preparations of individual seminiferous tubules to examine the interface between germ cells, Sertoli cells, and cells in the interstitial space. CSMD1 is expressed in a hatched pattern which is reminiscent of the actin bundles found at the Sertoli-Sertoli blood testis barrier and the Sertoli-spermatid interface (**Figure 2D**) (Lie et al., 2010).

**Figure 2.**
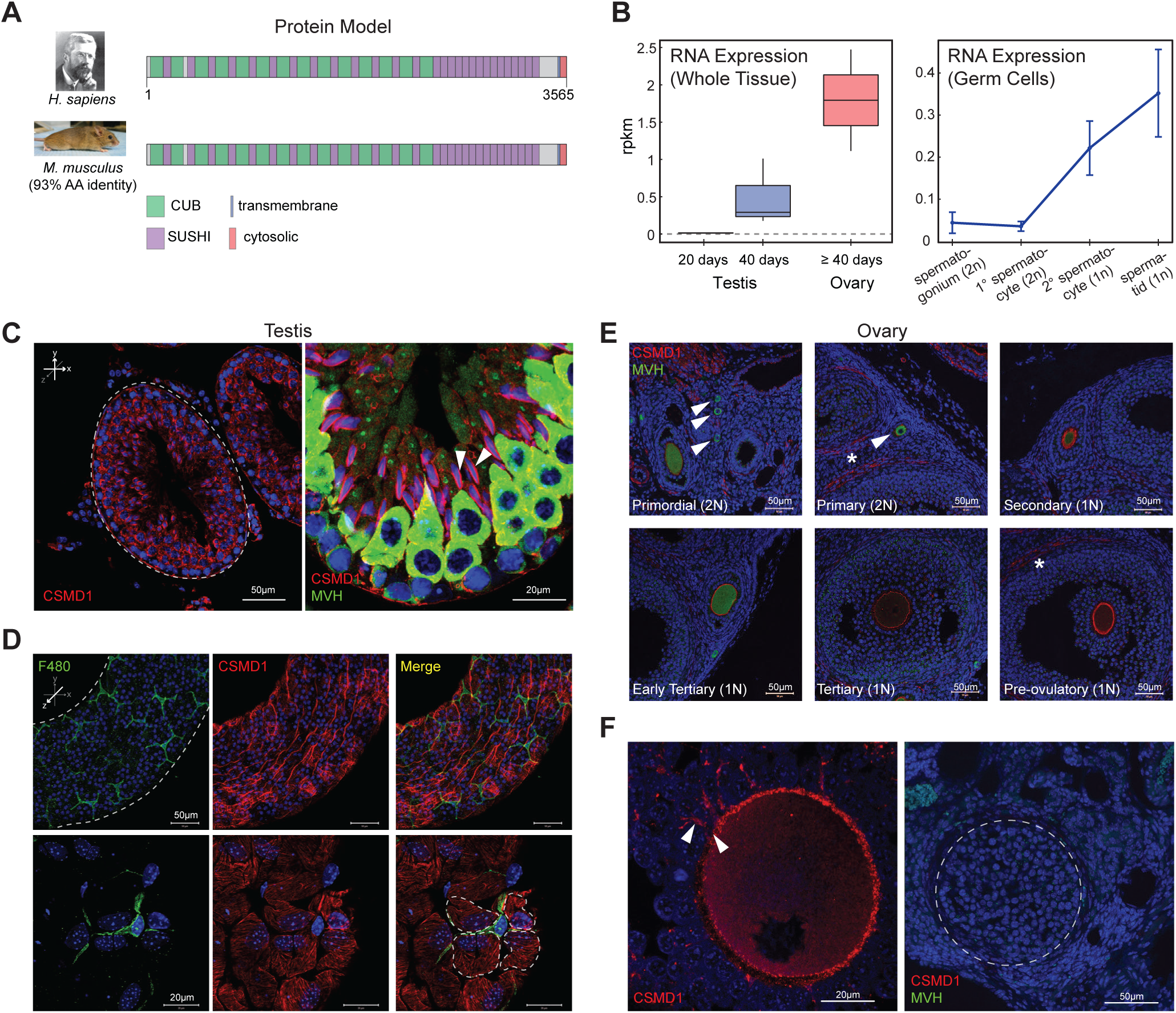
*Csmd1* is expressed in the male and female gonads. **(A)** Protein model of CSMD1 in human and mouse. CUB and Sushi domains, as well as the transmembrane and cytosolic domains are depicted along the protein model (97.1% of the CSMD1 protein is extracellular). (B) RNA expression of mouse *Csmd1* in sexually immature whole testes (20 days), sexually mature whole testes (40 days), and whole ovary. RNA-seq of FACS-purified germ cell populations show *Csmd1* expression changes during spermatogenesis. *Csmd1* RNA is maximally expressed at the spermatid stage of development. **(C)** Immunofluorescence (IF) visualization of CSMD1 (red) in testis seminiferous tubule cross sections (x-y axis). CSMD1 protein is broadly expressed in germ cells across all stages of spermatogenesis. MVH is a primordial germ cell marker whose expression peaks early, then steadily decreases during spermatogenesis/oogenesis. CSMD1 is maximally expressed on elongating spermatids during the spermiation process with somatic Sertoli cells (white arrowheads). **(D)** Whole mount testis tubule preparation (z axis). F480-positive macrophages (green) with characteristic ramified processes occupy the interstitial space. CSMD1 (red) is expressed in a hatched pattern which may correspond to the actin cytoskeleton of peritubular myoid cells and Sertoli cells. Four Sertoli cells surrounding a macrophage outlined by dotted lines. **(E)** IF of CSMD1 in developing oocytes (marked by MVH) and surrounding somatic cells. MVH expression decreases whereas CSMD1 expression increases with follicular development. CSMD1 is seen coating the oocytes at all follicle stages, occasionally staining weakly in granulosa cells, and staining more strongly on theca cells. Follicles named in each box are marked with white arrows when necessary. Theca cells are indicated by stars. **(F)** CSMD1 is maximally expressed at the oocyte surface and extends into the transzonal projections (white arrowheads), which physically connect the germ cell to the surrounding somatic granulosa cells. During ovulation the follicle releases the oocyte and regresses to form the corpus luteum (dashed lines). CSMD1 and MVH signal is absent.

Detailed analysis of the distribution of CSMD1 protein within the ovary revealed parallels with the testis. As in the testes, CSMD1 shows lower expression in follicles bearing diploid germ cells (i.e., primordial and primary follicles) and higher expression in follicles bearing haploid germ cells (i.e., secondary, tertiary, and pre-ovulatory follicles; **Figure 2E**). Theca cells also stain positive, as is quite apparent on late stage follicles (**Figure 2E**). The post-ovulatory corpus luteum shows no specific CSMD1 expression (dotted lines; **Figure 2F**). As with male germ cells, female oocytes require substantial physical interaction with surrounding somatic cells (Li and Albertini, 2013). At high magnification, CSMD1 is expressed along transzonal projections that emanate from the granulosa cells and connect to the oocyte membrane (**Figure 2F**).

### *Csmd1* knockout disrupts postnatal cellular development in multiple male and female tissues in mice

To confirm the biological role of *CSMD1* in male and/or female gonadal function, we perturbed its ortholog in a model organism. We generated a colony of *Csmd1* wildtype, heterozygous, and knockout mice and observed the effect of genotype on gonadal function and fertility (**Methods, Figure 3D**). In males, gross testis weight at necropsy did not differ significantly among wildtype, heterozygote, and knockout mice when measured in aggregate (P = 0.69). However, a subset of *Csmd1* knockout males suffer from profound anatomical and histological derangement of the testes (**Figure 3; Figure S3**). Remarkably, the most extreme instances of testes degeneration, Sertoli cell-only tubules, could be observed as early as 34 days of age (**Figure 3A; Figure S3B**). This time point corresponds to the onset of male sexual maturity (approximately 30-40 days) and the emergence of the spermatid germ cell stage, where *Csmd1* is maximally expressed. Males showed no evidence of derangement prior to sexual maturity (**Figure S3C**). Severity (“none”, “mild”, and “profound”) and onset (postnatal day 34 through day 300) of the degeneration phenotype vary greatly between individuals. In fact, different foci within the same testis of *Csmd1* knockout mice often show different stages of degeneration (**Figure 3B**). Our histological study of over 50 knockout animals uncovered two types of germ cell pathology whose connection to each other is unclear. The first is a sequence of active loss of germ cells within each tubule (**Figure 3B**). Spermatogenesis begins to become disorganized, especially at the late stages of spermiogenesis, with failure of spermiation, fewer numbers of elongating spermatids in the lumen, and mixing of spermatid steps in stages IX-XII. This is followed by the sloughing of all types of germ cells into the lumen; remaining germ cells can be observed in unusual tubules that appear to be missing one or more waves of spermatogenesis, and these eventually resolve as Sertoli cell-only tubules. Sloughed germ cells can be seen downstream in the epididymis, and, occasionally they obstruct the rete testis leading to dilation of the tubules (data not shown). These defects are most likely to arise due to disruption of interactions between Sertoli and germ cells. The second pathology was an apparent depletion of spermatogonial stem cells in the atrophic tubules; even in tubules with ongoing spermatogenesis, some areas show no spermatogonia. Significantly fewer germ cells express the male germ cell antigen TRA98+ (Poisson regression; P < 2 × 10^−16^; **Figures 3D and S3D**), in both atrophic and normal tubules, suggesting that knockout testes suffer from expression perturbations in addition to, or perhaps presaging, loss of spermatogonia and frank degeneration. Together, these observations indicate that the *Csmd1* knockout mutation (i) is not fully penetrant; and (ii) may be influenced by environmental and/or stochastic events. However, even after accounting for age covariates, *Csmd1* genotype segregates significantly with testes derangement status (P = 7.69 x 10^−3^; MANOVA; **Figure 3C**). Finally, we performed serial backcrossing for 9 generations on a subset of mice to validate the effect of the *Csmd1* null allele on a roughly constant genetic background (**Methods**). We recapitulated the degeneration phenotype in these backcrossed male knockouts (**Figure S3E**), indicating that *Csmd1* genotype status—not genetic background—was driving this signal of degeneration.

**Figure 3.**
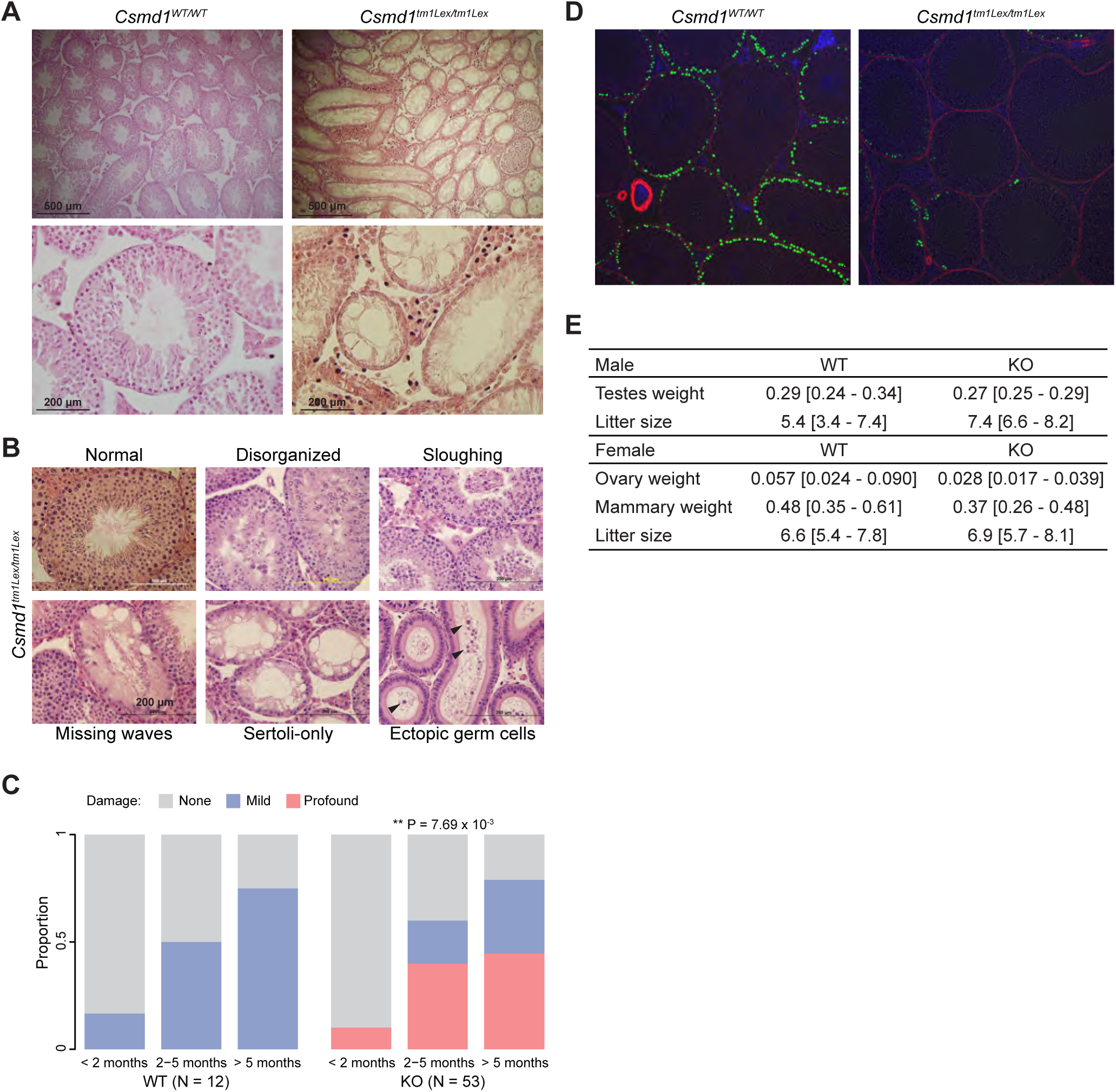
*Csmd1* knockout testes show profound and heterogeneous morphologic degeneration. **(A)** Seminferous tubule H&E histology of *Csmd1* wildtype and knockout littermates at 34 days of age. The majority of knockout tubules can be classified as “Sertoli cell-only” and contain no germ cells. Severe inflammatory changes are also observed in the interstitial space. **(B)** Qualitative stages of progressive morphologic degeneration. Seminiferous tubules from *Csmd1* knockout males showing normal morphology; loss of spatiotemporal architecture, but retained germ cells in all stages of spermatogenesis (“Disorganized”); loss of early stage germ cells into the tubule lumen (“Sloughing”); loss of all germ cells except late-stage spermatids (“Missing waves”); and loss of all germ cells, leaving a signature of vacuolization (“Sertoli-only”). Early-stage ectopic germ cells can be observed in the downstream epididymis, likely from upstream sloughing tubules. *Csmd1* knockout males can show multiple stages of degeneration within an individual testis. **(C)** Quantification of the degeneration phenotype. Testis H & E sections from control (n=12) and *Csmd1* knockout (n=53) animals were visually classified into one of three degeneration phenotypes: “None”, “Mild”, or “Profound” (Methods). The stacked barplots depict the proportion of damaged tubules among wildtypes and knockouts, stratified by age group. Damage severity segregates significantly with genotype, even after accounting for age (P = 7.69 x 10^−3^; ANOVA). **(D)** TRA98/GCNA positive spermatogonial cells (green) are much less abundant, and stain less intensely, in tubules of *Csmd1* knockouts. **(E)** Raw biometry and fecundity measurements from Csmd1 mutant colony. The mean and standard deviation of each measurement is reported. All weights are in grams.

In females, we observed severe inflammatory changes associated with foam cell infiltration, and, rarely, ovarian cysts in a subset of *Csmd1* knockouts (**Figures 4A and 4B**). Foam cells are multinucleated phagocytic macrophages which have become engorged with lipid, and are associated with ovarian aging. We performed Oil Red O staining which showed highly elevated lipid signal in the ovarian stroma of knockouts compared to age-matched controls, indicating a phenotype of premature ovarian aging in knockout animals (**Figure 4A**). *Csmd1* -deficient females had significantly smaller ovaries by mass when controlling for age, (p = 8.1 × 10^−3^; **Figure 3D; Figure 4C**). Furthermore, knockout females showed significantly more atretic follicles and fewer normal pre-ovulatory follicles at necropsy (p=3.5 × 10^−3^; Hotelling t-test; **Figure 4D**). To evaluate whether these biometric and histologic changes were also associated with reproductive performance, we estimated female time to pregnancy based on retrospective husbandry records. We generated a null distribution of time to conception which demonstrates distinct periodicity corresponding to the mouse female estrous cycle lasting 4-5 days (**Figure 4E**). Next, we stratified our population by maternal genotype. For *Csmd1* wildtype mothers, the bulk of conceptions occurred within the first estrous cycle as expected (Foldi et al., 2011), whereas most *Csmd1* knockout mothers became pregnant after two or more cycles (β_GT_ = 10.4; P = 0.012). A small minority of knockout females required many cycles to achieve pregnancy (> 60 days). Circulating gonadotropin levels did not differ between wildtype and knockouts after controlling for estrous stage, suggesting that this reduction in mating success was not secondary to impaired hormonal input along the HPG axis (**Methods, Figure S4**). Instead, if *Csmd1* knockout females bear a reduced ovarian reserve, there may be a reduced probability of conception per cycle due to a smaller oocyte target for male sperm. Interestingly, while knockout females achieved fewer pregnancies per estrous cycle, the average number of offspring born per pregnancy did not differ significantly between wildtype and knockout mothers (x̅_wt_ = 6.6 (95% CI [5.4-7.8]); x̅_ko_ = 6.9 (95% CI [5.7-8.1]); **Figure 3D**). However, pups borne of *Csmd1* knockout mothers suffered from significantly higher mortality rates during the neonatal period (1 - 10 days) when compared to wildtype/heterozygous mothers (% mortality_wT/het_ = 10.5% (95% CI [3.6% - 17.5%]); % mortality_KO_ = 50.0% (95% CI [30.0% - 70.0%]); Poisson regression P = 7.93 x 10^−7^; **Figure 5A**). We performed necropsy on expired offspring which revealed an absence of milk spots, suggesting death by starvation. Because neonatal mortality segregated with maternal genotype but not offspring genotype or paternal genotype, we hypothesized that this increase in mortality could be explained by a nursing defect in *Csmd1* -deficient mothers. Therefore, we performed IF to confirm that CSMD1 is expressed in the normal mammary gland through the adult life cycle of wildtype animals (**Figure 5B**). CSMD1 is observed on both luminal epithelial cells and myoepithelial cells of the mammary ducts, and on numerous stromal cells (**Figures 5B and 5C**). Ductal cell expression of CSMD1 appears to be regulated throughout the life cycle, with lowest expression seen in virgins, increasing in mid-pregnancy and lactation, with maximal expression during involution. Mammary glands from knockout females showed reduced density of the epithelial branching network during mid-pregnancy and post-nursing, likely explaining the lack of milk available to nursing pups (**Figure 5D**). Visual comparison of duct morphology in nulliparous wild type and knockout animals suggested that the main structural defect was a highly reduced incidence of lateral branches prior to pregnancy (**Figure 5E**), a conclusion that was statistically supported by quantitative image analysis (**Figure 5F**).

**Figure 4.**
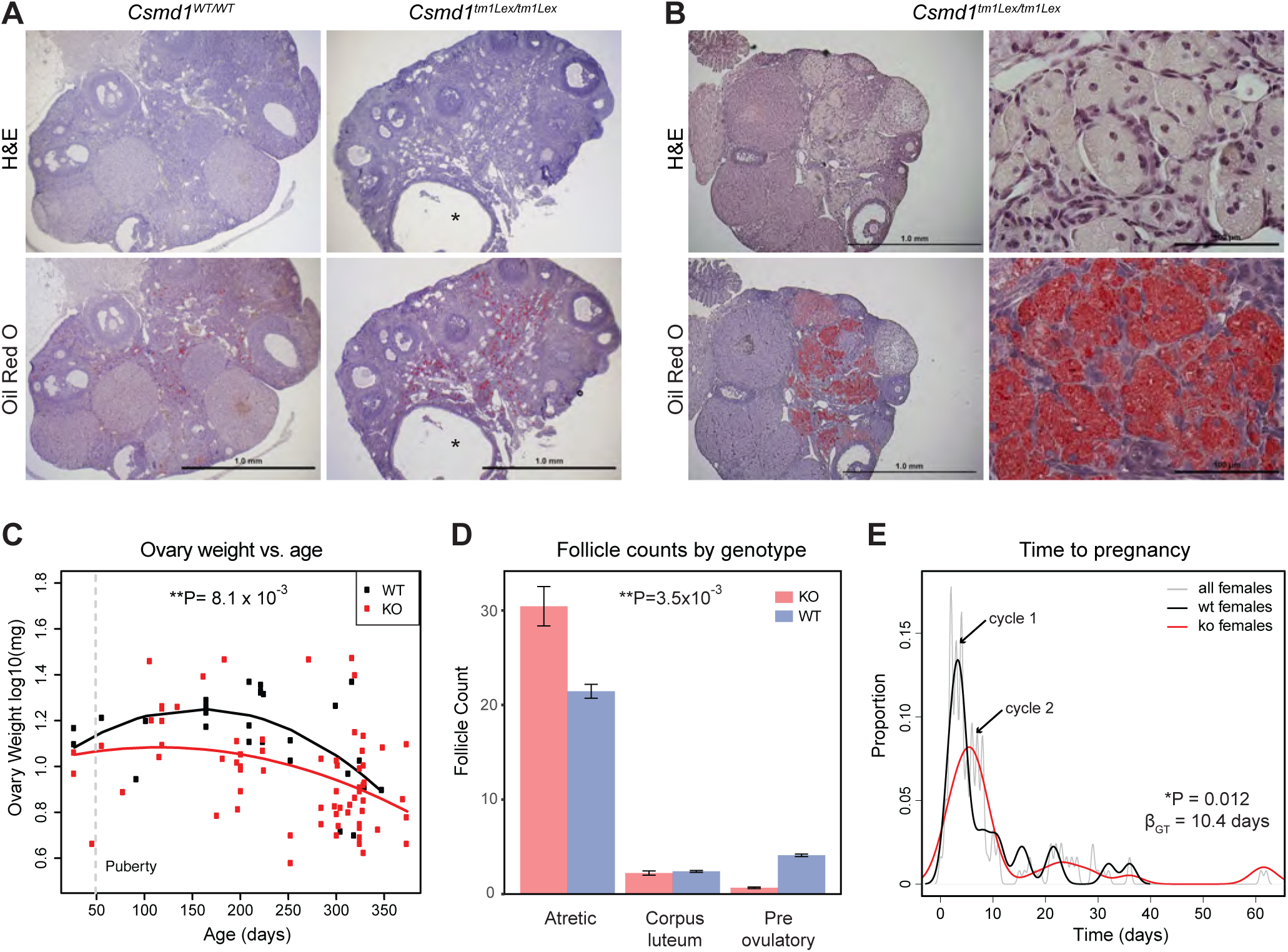
*Csmd1* knockout ovaries show reduced morphologic quality and reproductive performance. **(A)** Ovarian histology in *Csmd1* wildtype versus knockout females. Knockout ovaries were consistently enriched for foam cell macrophages, compared to age-matched controls, as detected by Oil Red O staining. Adjacent sections from 265 day old wild type (left)_and 240 day old knockout (right). Asterices indicate a large ovarian cyst in the knockout animal. Cysts were occasionally noted in knockout but not control animals. **(B)** Left: ovary from 336 day-old knockout showing extensive involvement of foam cells occupying 40% of the tissue. Right: high magnification of ovary section from same animal; top shows multinucleated appearance of foam cells, bottom is oil red O staining of same site in adjacent section. **(C)** Knockout females (n=68) have significantly smaller ovaries than wildtype (n=27) when controlling for age (p = 8.1 x 10^−3^). A quadratic regression model (shown) provided better fit to the data than a linear model. The grey hashed line indicates approximate onset of puberty in females. (D) Knockout females show more atretic and fewer morphologically normal pre-ovulatory follicles in ovary sections (p=3.5 × 10^−3^, Hotelling t-test). One section was evaluated per ovary. (E) Probability density plot depicting mating success over time, by female genotype. The probability density is periodic, corresponding to the female estrous cycle. Knockout females took significantly longer to achieve pregnancy (β_GT_ = 10.4, P = 0.01). All p-values reflect statistical models that account for confounders when appropriate such as age, body weight, and male factors.

**Figure 5.**
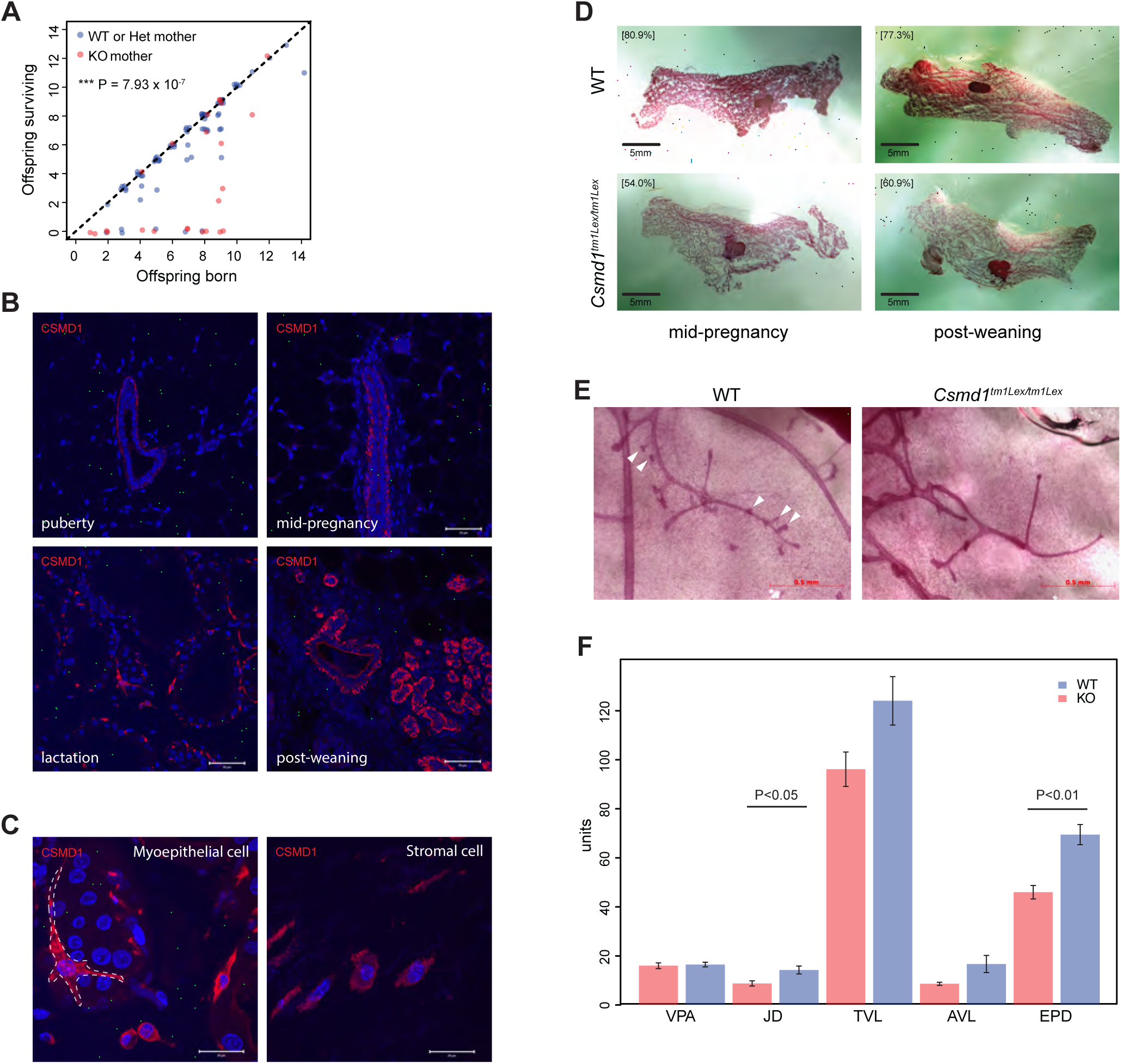
Knockout of *Csmd1* in mothers causes increased mortality in the offspring. **(A)** Scatterplot of number of pups surviving the neonatal period of 10 days (“Offspring surviving”) versus live births (“Offspring born”) versus, by maternal genotype (*Csmd1* wildtype and heterozygote versus *Csmd1* knockout). Points that lie along the dashed line (slope = 1) represent litters with no neonatal deaths. Maternal *Csmd1* genotype is significantly associated with surviving litter size (% mortality_WT/het_ = 10.5%; % mortality_KO_ = 50.0%; P = 7.93 × 10^−7^; Poisson regression). Data points deviate slightly from whole numbers for ease of visualization.**(B)** IF shows CSMD1 expression in bifurcating mammary ducts and bulbous terminal end buds. CSMD1 is expressed on both luminal epithelial cells and myoepithelial cells of the ducts throughout the adult life cycle: expression is lowest at puberty and increases during pregnancy, with the highest intensity during involution. Scale bars, 50 μm. **(C)** Close-up of CSMD1 on the membrane of a myoepithelial cell surrounding an alveolus during lactation (dashed line; left). CSMD1 was also observed on the membrane of small stromal cells (right). Scale bars, 20 μm. **(D)** Whole-mount mammary glands of *Csmd1* wildtype and knockout littermates during mid-pregnancy (left) and post-weaning (right). Square bracketed numbers represent normalized percent density of the branching epithelial network. Scale bars, 5 mm. **(E)** Knockout ducts appear to have greatly reduced lateral branching compared to wildtype (white arrowheads). Scale bars, 0.5mm. **(F)** To confirm this, we used computational image analysis to make quantitative comparisons of the structure and size of mammary ducts from age-matched nongravid nulliparous adults (n = 5 knockouts and n = 6 wildtype). Of 5 measurements made, two showed significant differences: the density of branch points along the duct (JD, p< 0.05) and the density of end segment (EPD, p < 0.01). VPA = percentage of area occupied by ducts. JD = density of branchpoints per mm. TVL = sum of Euclidean distances between all adjacent branch points. AVL = average length of Euclidean distance between adjacent branch points. EPD = number of duct end points normalized by total vessel length.

### The complement pathway is dysregulated in *Csmd1* knockout mice

The primary protein sequence of *CSMD1* shares homology with complement-interacting proteins (Kraus et al., 2006). Complement acts as an inflammatory/phagocytic signal in the innate immune system (Liszewski et al., 1996), and recent work has shown that classical complement components C1q and C3 are also responsible for microglia-mediated phagocytosis of excess neuronal cells in a normal developmental process known as synaptic pruning (Schafer et al., 2012). *CSMD1* (Schizophrenia Psychiatric Genome-Wide Association Study, 2011) (Schizophrenia Working Group of the Psychiatric Genomics, 2014) and complement *C4* (Sekar et al., 2016) have also been associated with schizophrenia in independent, well-powered human association studies. Furthermore, some of the most significantly associated variants previously associated with azoospermia encompass the greater MHC locus, which include complements *C2, C4* and factor B (Ni et al., 2015; Zhao et al., 2012). *Csmd1* is also known to inhibit the classical complement pathway *in vitro* (Escudero-Esparza et al., 2013; Kraus et al., 2006). Thus, to consolidate the putative roles of complement with *Csmd1*-mediated pathology, we investigated the activity of macrophages and complement component C3 in wildtype and *Csmd1-null* gonads. *C3* mRNA is detectable in whole testes and ovaries, and in testicular germ cells at multiple stages of spermatogenesis (**Figure 6A**). *C3* and *Csmd1* mRNA expression are anticorrelated throughout spermatogenesis. Macrophages, the immune cells most commonly associated with complement-mediated phagocytosis, are found in the interstitial space between seminiferous tubules (**Figure 6B**). We frequently observed C3 in the interstitial space, but not within the tubules; likewise, C3 could be observed further downstream in the epididymis, in the peritubular regions but not inside the lumen (**Figure 6B**). We measured bulk macrophage content and complement C3 deposition in *Csmd1* wildtype and knockout testes (**Figure 6C; Figure S5; Methods**). The proportion of C3-positive cells is significantly higher in *Csmd1* knockout versus wildtype testes (x̅_wt_ = 0.017; x̅_ko_ = 0.066; ANOVA P = 7.7 × 10^−4^), consistent with an inhibitory role for *Csmd1* against complement.

**Figure 6.**
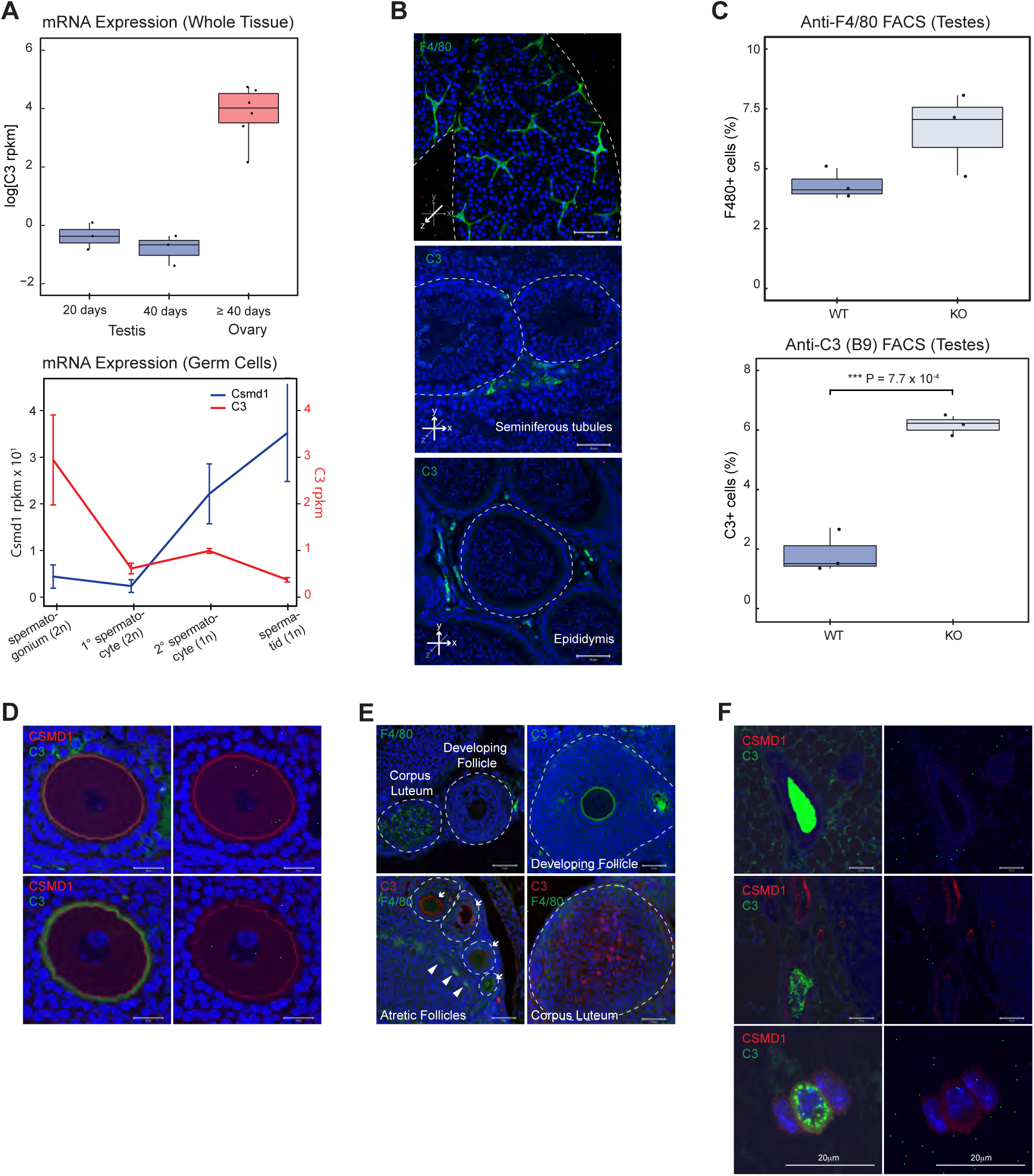
Complement C3 regulation in mouse testes and ovaries. **(A)** RNA expression of mouse *C3* in sexually immature whole testes (20 days), sexually mature whole testes (40 days), and whole ovary. RNA-seq of FACS-purified germ cell populations show *C3* expression changes during spermatogenesis. *C3* is maximally expressed at the spermatogonium stage. *Csmd1* RNA-seq data from Figure 2 are rescaled and superimposed for ease of comparison. **(B)** Complement and macrophages are confined to the basal compartment in normal tubules. Whole mount tubule IF visualization of macrophage marker F480 along shows extensive macrophage abundance along the interstitium. Cross section of tubule shows C3 expression in the interstitium, but not within the lumen. Cross section of downstream epididymis also shows continued exclusion of C3 from the lumen. Individual tubules are circumscribed by dashed lines. **(C)** Boxplots depicting F4/80 abundance and C3 deposition in FACS-sorted *Csmd1* wildtype versus knockout testes. Both F4/80 and C3 are increased in knockout testes, though only significant for C3 (2-tailed t-test; P = 7.7 × 10^−4^). **(D)** C3 and CSMD1 co-localize at the oocyte surface. In most follicles examined, C3 and CSMD1 co-localize at the oocyte plasma membrane with overlapping signal (top 2 panels). On rare occasion the two signals separate and C3 stains in a ring outside of CSMD1 (bottom 2 panels). **(E)** F4/80 IF of adjacent follicles shows positive signal in corpus luteum, but not in follicles, consistent with prior expectations. C3 is expressed on the oocyte as well as in the follicular fluid of the developing antrum (asterisk). Atretic follicles at different stages of degeneration (white arrows) show varying levels of C3 and F4/80 expression. F4/80 can also be seen in a punctuated pattern along the stroma and thecal layers (white arrowheads), consistent with prior expectations. C3 signal is also seen in corpus luteum. Individual follicles are circumscribed by dashed lines. (F) C3 and CSMD1 also colocalize in mammaries. As early as puberty there is abundant C3 staining within mammary ducts with empty lumens (top panel) and ducts with cells in the lumen (middle panel). C3 is also present in vesicles of some CSMD1+ stromal cells (bottom panel).

In wildtype ovaries, we observed a localization of C3 and macrophages that support the hypothesis that complement-mediated phagocytosis and cellular remodeling are processes that regulate normal gonadal function. Interestingly, C3 is localized to the oocyte surface in normal developing follicles, colocalized with CSMD1, and then observed to be diffused in large amounts throughout the corpus luteum, which is devoid of CSMD1 (**Figures 6D and 6E**). Macrophages are a prominent cell type in the ovary and associated with, but excluded from entering, healthy follicles; they invade corpora lutea and degrading follicles (**Figure 6E**). As predicted in a model of C3-mediated phagocytosis by macrophages, C3 colocalizes with macrophages in the corpus luteum as well as in atretic follicles. Interestingly C3 is abundant within the early follicular antrum (probably in follicular fluid), suggesting that C3 may be important for remodeling the connections between granulosa cells during antrum formation (**Figure 6E**). It has previously been reported that activated C3 is present in human follicular fluid at levels comparable to sera, but its physiological role in folliculogenesis, ovulation or fertilization is unknown (Perricone et al., 1992).

Finally, we observed a pattern of C3 and CSMD1 expression in wildtype mammaries that also supports the notion that CSMD1-complement interactions are dysregulated in the pathologies observed in CSMD1 knockouts (**Figure 6F**). As early as puberty, C3 can be seen in high levels within the mammary duct lumen of virgin animals. We speculate that C3 may be involved in the process of lumen formation, as lower levels of C3 are observed in lumens that are just beginning to open and contain dissociated cells. C3 is also expressed within vesicles of specific subsets of CSMD1-positive stromal cells, likely macrophages or eosinophils.

Based on previous findings that CSMD1 is a negative regulator of C3, we predicted that removal of C3 would partially or completely alleviate the morphological degeneration and fertility defects observed in *Csmd1* knockout mice. To test this prediction, we generated a colony of *C3/Csmd1* double knockout (DKO) mice. Surprisingly, we found no evidence of rescue in DKO males or females (**Figure S6**). Instead, we observed an unmasked phenotype of more severe histological degeneration in all DKO females, characterized by even more invasion of foam cell macrophages, extensive pyknosis, and deformed follicles. We also observed profound inflammatory changes in the mucosal layer of the oviduct (**Figure S6B**). We monitored the fertility of 19 DKOs (10 males and 9 females), and of these, only 4 (21%) produced progeny after at least 3-7 months of mating (3 males and 1 female; **Table S3, Figure S6C**). The average litter size resulting from successful mating was small compared to wildtype (mean size 4.25 pups). These extreme phenotypes are not observed in *Csmd1* nor *C3* single knockouts, indicating that the combined effect of *Csmd1* and *C3* on fertility is synergistic.

## Discussion

We used a human genetic screening approach to identify genes that modulate male and female gonadal function, and identified a strong candidate, the complement regulator *CSMD1.* The human phenotypes that we studied were ascertained for having abnormal, early loss of germ cell development, and we observed defects in gametogenesis in both male and female *Csmd1* knockout (KO) mice. We performed a series of experiments with mice to evaluate three competing explanations for this germ cell loss: increased cell death, failure of proliferation, or increased phagocytosis.

During our work-up of testis pathology in sexually mature *Csmd1*-null mice, we observed neither qualitative nor quantitative differences in the abundance of apoptosis markers TUNEL in testis cross sections or Annexin-V in dissociated whole testis FACS (data not shown). These observations, coupled with adult onset of the testes degeneration phenotype do not support an increase in apoptosis as the mechanism for gonadal dysfunction.

Because much of the hormonal and cellular machinery for cell division is shared between both sexes, a failure of proliferation either due to endocrine disruption or maturation arrest is another possible explanation for infertility in males and females. We excluded systemic endocrine defects that would be observed in the case of failure of the hypothalamus or pituitary (**Figure S4**). We did not observe any stage-specific accumulation or depletion of germ cells in either sex, nor, as mentioned above, any tell-tale signs of excess apoptosis that is usually seen in such cases (Lipkin et al., 2002; Yatsenko et al., 2015). We observed no significant differences in PCNA marker levels between *Csmd1* wildtype and knockout testes of adult animals (data not shown).

The finding of increased C3 deposition in testis, coupled to the known complement-regulatory function of CSMD1, suggests that improper phagocytosis of cells or cellular structures underlies at least part of the defect, but does not illuminate the cell type(s) or biological process(es) that are consequently affected. The diverse defects observed by histology points to a problem in maintenance of the stem cell niche or stem cell function, and possibly Sertoli cell function. We see no consistent signs of defects in germ cell morphology or stage-specific depletion or enrichment of cells. There were no overt signs of derangement of Sertoli cell phagocytosis, such as a universal bloating of Sertoli cell vacuoles, or the abnormal presence of elongated spermatid heads near the basement membrane, in all *Csmd1* KOs investigated. We observed no evidence of complement deposition inside the lumen of the seminiferous tubules. It is widely believed that the tubules are an immune privileged site, and we observed no evidence of macrophages inside the tubules or disruption of the blood-testis-barrier (BTB) in knockouts. However, both spermatogonial stem cells and Sertoli cells exist outside the BTB, and macrophages have been shown to be required for proper SSC differentiation (DeFalco et al., 2015). In further support of a niche defect, the strongest quantitative difference in protein abundance observed between wildtype and *Csmd1* KO, among over 20 proteins tested, was a universally lower expression of the germ cell nuclear antigen TRA98 in spermatogonia (**Figures 3D and S3D**). Our results are also consistent with a role for Sertoli cells in the pathology of knockouts, either due to interactions with germ cells or interstitial cells, as we see extensive sloughing of germ cells from the epithelium, as well as rare whorls of Sertoli cells in the lumen (**Figure 3B**).

The histology data from ovaries are consistent with a model where dysregulation of the macrophage-complement axis leads to loss of developing follicles and/or oocytes. Macrophage activity in the ovary is very carefully regulated in time and space during the estrous cycle. It is well known that macrophages are physically associated with most if not all developing follicles, and that this association is not just a response to atresia (Gaytan et al., 1998; Tingen et al., 2011). After ovulation, macrophages invade the ruptured follicle which undergoes apoptosis/phagocytic luteolysis, forming the corpus luteum (Kato et al., 2005). CSMD1 is also highly expressed on oocytes of the developing follicle, but not in the corpus luteum (**Figure 2E, F**). Disruption of CSMD1 function may allow for premature macrophage invasion of the developing follicle, leading to excessive oocyte atresia, fewer ovulations, and reduced probability of pregnancy (**Figure 4D,E**). *Csmd1* null females give birth to normal litter sizes, which limits the possibility that CSMD1 mediated follicle loss occurs during the cyclic recruitment of antral follicles. Instead it may be operating at the phase of initial recruitment, or perhaps even earlier in the establishment of the oocyte or follicle pool. A more sensitive analysis of oocyte and follicle counts at multiple time points will be needed to pinpoint exactly where in the oocyte lifecycle atresia occurs.

In addition to the gonads, *CSMD1* also governs post-natal developmental processes across other tissue systems. We have demonstrated a robust association between neonatal mortality rate and maternal *Csmd1* genotype status, with corresponding reduction in the epithelial network of the maternal mammary gland (**Figure 5**). The mammary gland is a highly motile network of branching epithelial tissue that advances and recedes during different stages of post-natal development (i.e., puberty, pregnancy, nursing, etc. (Sternlicht, 2006)). The characteristic directionality of mammary branching is conferred by polarized cell proliferation and phagocytosis mediated by macrophage remodeling, especially in anticipation of nursing (Pollard, 2009). Furthermore, multiple complement and complement-regulatory components are robustly upregulated during periods of apoptosis and phagocytosis in the mammary tissue of multiple species including humans (Clarkson et al., 2004; Laufer et al., 1999), though the functional significance of this regulatory pattern is unknown. Breast milk itself also suppresses complement activation (Ogundele, 1999). Finally, CSMD1 is expressed on the luminal aspect of mammary ducts and terminal end buds, where much of the pregnancy-associated breast remodeling occurs (**Figure 5B**) (Kamal et al., 2010; Kraus et al., 2006). We show a reduction in mammary epithelial density, due to reduction in secondary or tertiary branch points, whose normal geneses are governed by multiple up- and down-regulatory chemotactic signals in concert with physical interaction with phagocytic immune cells (i.e., macrophages)(Ingman et al., 2006)

It is possible to propose a model to reconcile the pathology that we observe across multiple tissues in *Csmd1* null animals. While complement has a well-appreciated role in innate immunity, evidence is beginning to emerge that it plays a role in the regulation of self cells during normal human development. A key example of this biology is the recent description of complement-mediated synaptic pruning in the developing brain, a process that is under genetic influence and can confer risk for disease when dysregulated (Schafer et al., 2012; Sekar et al., 2016). Here, we have reported complement-associated pathology of post-natal developmental processes in three additional tissues in *Csmd1* null animals; we have observed complement protein expression in all three tissues, and macrophages have been shown to be essential for normal development in all three tissues. A parsimonious model to describe the set of defects we observe here is that macrophages (and perhaps other phagocytes) regulate and refine developing cells in testis, ovaries, and mammary by controlled deposition of complement onto their cell surface.

In this model, differentiating cells that progress through developmental checkpoints upregulate complement regulators on their cell surface. Healthy, well-formed cells (including macrophages) secrete C3 at low level continuously into interstitial space and possibly onto the surface of cells. A function of this local complement synthesis is low grade activation to get rid of “junk,” without an adaptive immune response or very vigorous innate one. Intracellular and extracellular C3 is available to tag and mark unwanted cells or cell-derived structures for removal. CSMD1 (and presumably other complement regulators) has a special function that involves complement modulation at a highly localized and specific immune privileged site. A certain amount of activated C3 fragments need to be deposited on a target to carry out a specific function – too much or too little has a “bad” consequence. We call this controlled phenomenon targeted and restricted activation of the complement system (TRACS), and initially described the concept of TRACS to explain the function of the complement regulator membrane cofactor protein (MCP) in controlling complement deposition on the inner acrosomal membrane of acrosome-reacted spermatozoa (Riley-Vargas et al., 2005).

In our model, there is normally limited or no engagement of adaptive immune players who are not present near the site of TRACS and require more of an acute inflammatory setting to get there. CSMD1 deficiency may be enough to periodically tip the balance of controlled complement activation towards a pathogenic outcome. Conversely, if debris is not removed, developmental processes are disorganized or blocked. In this way, C3 inactivation is predicted to exacerbate, not rescue, the fertility defects we observed in *CSMD1* null animals. If complement marks targets of phagocytosis in the testis as previously shown in the brain, ectopic complement expression across the BTB may inappropriately activate the apoptotic and phagocytic apparatus in *Csmd1* KO testes. Remarkably, *TEP1* (a distant ortholog of C3) has been shown to clear apoptotic germ cells in the mosquito testis by this very process (Pompon and Levashina, 2015) .

The TRACS model is consistent with known molecular functions of macrophages and complement. However, macrophages have recently been shown to regulate the spermatogonial stem niche by an unknown molecular mechanism (DeFalco et al., 2015). The defects we observe in CSMD1 -/- males are consistent with a niche problem, and we speculate that controlled complement deposition on spermatogonial cells could mediate interactions with macrophages.

The unmasking of a more severe phenotype in *C3/Csmd1* DKO mice is an unexpected but previously documented signature of complement-mediated disease. For example, double knockout of complement factor H (*CFH*) and factor P (*CFP*) unexpectedly converts mild C3 glomerulonephritis to lethal C3 glomerulonephritis in mice(Lesher et al., 2013). Similarly *CFH/C3* DKO unexpectedly unmasks a more severe form of age-related macular degeneration in mice(Hoh Kam et al., 2013). Multiple explanations for this phenomenon have been set forth, including a dual role of C3, differences between fluid-phase and local C3 activation, and C3 gain of function. More extensive mutation constructs including conditional knockouts and allelic series may help to distinguish among these scenarios.

Finally, we note that our observations may be informative about the biological basis for the highly elevated mutation rate over *CSMD1.* Recently, it has been reported that the two hottest hotspots for maternally derived DNMs in humans are centered on two large genes: *CSMD1* and *WWOX* (Goldmann et al., 2016). These two genes are also among the top 27 most frequent sites of double-strand break formation in primary neural progenitor cells(Wei et al., 2016). Careful study of CNV mutation mechanisms has led to a specific model for the genesis of CNVs over large genes, known as Transcription-dependent Double-Fork Failure (TrDoff), whereby transcription of large genes interferes with DNA replication (Wilson et al., 2015). The TrDoff model predicts that duplications will be enriched at the edges of large gene, while deletions are enriched in the gene body, a pattern that is consistent with our data on *CSMD1* (**Figure 1B**). We have observed that CSMD1 protein is present in primordial follicles of adult mice, suggesting that *CSMD1* is transcribed in oocytes throughout most, perhaps all, of the life of the animal. We predict that this constant, sustained transcription of a large gene in each oocyte may expose the female germline to transcription-coupled molecular conflicts like TrDoff that are not as pronounced in the male germline. We speculate that this could be exacerbated by incomplete DNA replication at *Csmd1* at the time that the oocytes arrest in MI, and differences in the amount of replication stress experienced during the initial expansion of the germ cell pool, which happens more quickly in females compared to males. We predict that the *WWOX* is also expressed in oocytes with a developmental timing similar to *CSMD1.*

In conclusion, we have used human genetics and animal models to identify a likely role for the complement system in postnatal developmental processes across multiple tissues in the body. When combined with existing observations from mammalian brain and other model organisms, we predict that macrophage mediated complement activity on self cells is a normal and highly controlled process in many developmental systems in metazoans. Our work highlights the need for deeper investigation into the role of immune system components in reproductive tissues, and the opportunities that such work can have to illuminate and connect common biological processes that produce disease in more complex contexts across the body.

## Materials and Methods

### Human Patient Populations

We used male infertility case-controls cohorts that were previously described (Huang et al., 2015; Lopes et al., 2013).

#### WHI-SHARe

To create an analogous case-control cohort of female gonadal dysfunction, we turned to the SNP Health Association Resource (SHARe) cohort studied under the umbrella of the Women’s Health Initiative (WHI)(Hays et al., 2003). We constructed POI case and control definitions from the dense reproductive phenotype data collected on each subject. A self-reported age of menopause before 40 years was used as the only case inclusion criterion. Case exclusion criteria were oophorectomy prior to age 40, a diagnosis of lupus or rheumatoid disease, and a “yes” answer to the question “Did a doctor ever say that you had cancer, a malignant growth, or tumor?”. Smoking history, which is a known factor influencing ovarian reserve, was controlled for during the analysis of genetic data.

#### UK Biobank

We generated a table of phenotype data for constructing POI case and control labels using controlled-access data from the UK Biobank. Exclusion criteria for the study were: withdrawn consent, poor heterozygosity or missingness as defined by the UK Biobank; > 10 relatives in the UK Biobank cohort; not used in autosome phasing, apparent sex chromosome aneuploidy; mismatch between genetic and self-reported sex; ever smoker; any self-reported non-Caucasian ancestry; prior diagnosis of rheumatoid arthritis, lupus, or pelvic cancer; mismatch between self-reported ethnicity or age at menopause among three assessments; SNP array call rate <98%. In the case of pairs of 2^nd^ degree relatives or closer, the one individual with the lower SNP-call rate was dropped. The inclusion criteria for POI case status were self-reported age of menopause < 40 years old, and all remaining individuals in the cohort (after exclusions mentioned above) were used as controls.

#### Parent-Offspring Trios

For estimation of chromosome 8 human *de novo* mutation rates, blood samples were collected from parent-offspring trios, parent-twin quartets, and parent-triplet quintets who delivered at Inova Fairfax Hospital and whole genome sequence data were acquired as part of the Inova Translational Medicine Institute’s Premature Birth Study as described previously (Goldmann et al., 2016).

### Mouse colony breeding

We acquired a constitutive *Csmd1* knockout mouse (*Mus musculus*) on a mixed 129SvEvBrd:C57BL/6 background from the UC Davis KOMP Repository (Project ID CSD118901). The original construction of the mouse is described previously(Friddle et al., 2003). Briefly, a 1.086 kb deletion encompassing *Csmd1* exon 1 and part of intron 1 were replaced with a lacZ/neomycin cassette. Deletion of this segment was confirmed with Southern blot and PCR. Due to the extreme size of *Csmd1* (1.6 Mb), we also analyzed RNA seq data across all 70 exons in knockout testes and ovaries. In ovaries, knockout read counts relative to wildtype are suppressed across all 70 exons. In testes, knockout read counts relative to wildtype are broadly suppressed across exons 1-57 and upregulated from exons 58-70. The amino acid coding portion of these upregulated exons range in size from 45 bp to 180 bp. The translational viability of these fragments is unknown. All littermate tissue comparisons in this study (described below) were generated from dam_heterozygous_ x sire_heterozygous_ crossings from this original colony. Next, to eliminate variance in phenotype explained by variance in background genotype (if any), we serially backcrossed the *Csmd1* mutation onto a constant C57BL/6 background for 5 generations. From this F5 backcross generation we performed a dam_heterozygous_ x sire_heterozygous_ cross from this to create wildtype and knockout littermates, and performed analogous histology and immunofluorescence experiments as with the original colony (described below; **Figure S7**). We performed microsatellite genotyping of these littermates to estimate the C57BL/6 background after backcrossing (Washington University Rheumatic Disease Core). We estimated the F5 proportion of C57BL/6 ancestry of 0.91 (95% CI [0.89-0.93]). For DKO experiments, we introgressed a *C3* mutant line described previously (Circolo et al., 1999) until we achieved *Csmd1/C3* DKO mice.

### CNV and SNV discovery

Array data for the Women’s Health Initiative SHARe cohort were downloaded from the NCBI Database of Genotypes and Phenotypes (dbGAP accession number phg0000g1.v2). SHARe samples were genotyped on the Affymetrix 6.0 platform. We created a high-quality set of CNV calls for all cohorts using our own internal pipelines. SHARe samples were processed with Affy6CNV (a wrapper that we wrote for the Birdsuite package (Korn et al., 2008)) for data processing and QC. We obtained raw SNP array data from the UK Biobank and performed single sample CNV discovery using PennCNV(Wang et al., 2007). Individuals with > 200 CNV calls were dropped. CNV calls with PennCNV quality score > 30 were retained, and adjacent CNVs in the same sample were merged.

Exome sequencing was performed on a subset of the WHI subjects as part of the Women’s Health Initiative Sequencing Project (WHISP); all available WHISP BAM files were downloaded from the NCBI Database of Genotypes and Phenotypes (dbGaP accession phs000200.v10.p3.c1 and phs000200.v10.p3.c2)(Tryka et al., 2014). Genotypes were recalled, jointly, from 1,668 WHI BAM files using Haplotype Caller, recalibrated and cleaned according to GATK best practices using GATK-3.2.2.

### Association testing

#### CNVs

Rare CNVs were associated with case-control status using generalized linear models. For the SHARe association analysis, we included the top 10 ancestry eigenvectors (calculated from the full SHARe genotype matrix) and smoking history as covariates. For the UK BioBank association analysis, individuals with any history of smoking were excluded from the analysis, and we included as covariates BMI and the top 10 ancestry eigenvectors calculated from the full UK Biobank SNP genotype matrix.

#### SNVs

We tested for an association between rare SNVs in *CSMD1* and age at menopause in the WHI samples using the Sequence Kernel Association Test (SKAT) (Lee et al., 2012), weighting each variant with the Combined Annotation Dependent Depletion (CADD) value (Kircher et al., 2014). Five ancestry eigenvectors and smoking history were included as covariates; significance was evaluated using bootstrapping with 5,000 samples.

### Testes dissociation, cell sorting, and RNA extraction

Sexually mature (40 ± 1 days old), wildtype male mice were sacrificed, and their testes were decapsulated and homogenized in a 1X MEM solution (Gibco 11430-030) containing 120 U/mL Type I Collagenase (Worthington Biochemical LS004194) and 1 mg/mL DNAse I (Roche 10104159001), and agitated for 15 minutes. 1X MEM was replaced and added with 50 mg/mL Trypsin (Worthington Biochemical 54J15037) and 1 mg/mL DNAse I and agitated for 15 minutes, then mechanically homogenised for 3 minutes. 50 mg/mL Trypsin and 1 mg/mL DNAse I were added and agitated again for 15 minutes. We added 0.4 mL heat inactivated Fetal Bovine Serum (Sigma F1051), 5 μL Hoescht 33342 (Life Technologies H3570), and 1 mg/mL DNAse I, and agitated for 15 minutes. Individual cells were dissociated by pipetting sequentially through two 40 μm cell strainers (Falcon 352340). For each individual mouse, one dissociated testis was used for wholetissue RNA extraction and sequencing, and the other testis was used for germ cell purification, RNA extraction, and sequencing. All dissociation steps were performed at 33°C. Dissociated testes were sorted as described previously on a modified MoFlo cytometer (Beckman Coulter) at the Washington University Siteman Flow Cytometry Core using a krypton-ion laser (Lima et al., 2016). Cells that are stained with Hoechst can be clustered in two wavelengths: (i) blue fluorescence, which is informative of DNA content, and (ii) red fluorescence, which is informative about chromatin state and Hoechst efflux from the cell. Based on these parameters, we separated homogenised testes suspensions into four purified populations: (i) spermatogonia, (ii) primary spermatocytes, (iii) secondary spermatocytes, and (iv) spermatids. These separated populations were collected and RNA extraction performed on them. RNA from whole testes was extracted with the RNeasy Plus Mini Kit (Qiagen 74134), and RNA from FACS-purified germ cell populations was extracted with the RNeasy Plus Micro Kit (Qiagen 74034).

### RNA-seq

Whole testis, whole ovaries, and purified male germ cell subpopulations were obtained from wildtype and *Csmd1* null siblings. We extracted polyadenylated mRNAs from each tissue/cell type and converted these into RNA-seq libraries. Three biological replicates of each tissue or cell type were sequenced with a 2 x 101bp paired-end protocol. Reads were mapped to Ensembl Mus musculus reference R72 and transcript expression levels were summarized as reads-per-kb of exon per million-mapped reads (RPKM) using the TopHat2 package(Kim et al., 2013). RPKMs were adjusted for batch effects and cryptic covariates using PEER(Stegle et al., 2012), quantile normalized, and then the R package poissonSeq was used for differential expression analyses(Li et al., 2012).

### Immunostaining and imaging

Testes and ovaries were dissected, fixed in 4% paraformaldehyde (Electron Microscopy Sciences), and embedded in paraffin. We baked 5μm sections at 60°C for 1 hr, deparaffinized in Xylenes, and rehydrated into PBS (Corning). Antigen retrieval was done in boiling citrate buffer (10mM sodium citrate, 0.05% Tween-20, pH 6.0) for 20 min. Sections were blocked in PBS containing 0.2% Triton X-100 and 5% normal donkey serum (Jackson Laboratories) for 1 hr at room temperature and then incubated with primary antibodies diluted in blocking solution over night at 4°C. After washing with PBS-Tx (PBS containing 0.2% Triton X-100), they were incubated with fluorescent secondary antibodies in blocking solution for 1 hr at room temperature, washed with PBS-Tx, and treated with 0.2% Sudan Black in 70% EtOH for 10 min, followed by PBS washes. The sections were then counterstained with Hoechst dye 33342 diluted 1:500 in PBS for 5 min, washed once with PBS-Tx for 2 min and then with PBS, and mounted in ProLong Diamond anti-fade mounting medium (Molecular Probes). Imaging was done on an Olympus LSM700 confocal microscope using Zen software, and images were processed using Photoshop CS5 (Adobe). Antibodies used were gt α-CSMD1 N20 (Santa Cruz Biotechnology, 1:100), rb α-mouse vasa homolog (MVH) (Abcam 13840, 1:1,000), donkey α −gt CF594 (Biotium, 1:300), and donkey α -rb Alexa488 (Life Technologies, 1:300), rb α- β-gal (Cappel 1:333), rat F4/80 BM8 (Santa Cruz Biotechnology 1:50), donkey α -rat Alexa488 (Life Technologies, 1:300), rb α-C3 (Abcam 200999, 1:2,000), and gt α-rb Alexa568. Whole mount IF samples were prepared as described previously (DeFalco et al., 2015). For immunohistochemistry, 5μm paraffin sections were treated as above, except the secondary antibody was biotin-coupled horse a-goat (Vector Laboratories, BA-9500, 1:200), and detection was done using the Vectastain Elite ABC kit (Vector Laboratories, PK-6100) and DAB Peroxidase Substrate kit (Vector Laboratories SK-4100) per the manufacturer’s instructions. Sections were counterstained with hematoxylin, mounted in Cytoseal Xyl (Thermo Scientific), and imaged on a Zeiss Axioplan 2 microscope equipped with an Olympus DP71 camera and DP software. X-gal staining was performed as described previously (Li et al., 1998) .

### Histology

Freshly-dissected gonads were fixed under agitation in Modified Davidson’s fixative (Electron Microscopy Sciences 64133-50) for 24 hour and Bouin’s fixative (Electron Microscopy Sciences 26367-01) for 24 hours. Fixed tissues were embedded in paraffin and sectioned at 5 μm. Sectioned tissues were stained with hematoxylin and counter-stained with eosin. Stained testes from 65 individual mice of known age and genotype (12 wildtype, 53 knockout) were provided to a single mouse pathologist in a blinded fashion. All samples received a score of 0 (no damage), 1 (mild damage), or 2 (profound damage) (see **Figure S3A** for examples). In order to estimate the effect of genotype on score, we fit a linear analysis of variance model:

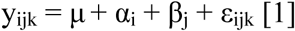

where y_ijk_ is the damage score for individual *k*, μ average damage score across all animals, α_i_ is the effect of genotype *i*, β_j_ is the effect of age *j*, and ε_ijk_ is the random error associated with the *k*th observation.

### Germ cell quantification

We performed immunofluorescence as described above on a pair of 34 day old male littermates (the same individuals as seen in **Figure 3A**) using anti-TRA98 antibody (Abcam ab82527). We generated count data for total cells (filtering based on size and shape), and for TRA98-positive cells (filtering based on green fluorescence) using the ImageJ software package. In order to estimate the effect of genotype on TRA98 cell count, we fit the following linear model:

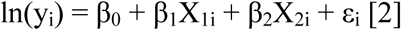

Where y_i_ is the TRA98-positive count in image *i*, and X_1_ is the genotype (*Csmd1* wildtype versus knockout), and X_2_ is the total cell count. ε_i_ is the nuisance variable for image *i*.

### Gonad size analysis

We sacrificed 229 adult mice (106 males and 123 females), and measured body weights and bilateral gonad weights at necropsy. For males, mean body weight was 37.1g, mean testes weight was 273mg, and mean age was 201 days. For females, mean body weight was 31.3g, mean ovary weight was 32mg, and mean age was 234 days. In order to estimate the effect of genotype on gonad weight, we fit the following linear model:

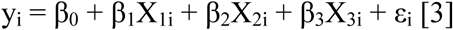

Where y_i_ is the gonad weight in individual *i*, and X_1i_, X_2i_, and X_3i_ are the genotype, age, and body weight of individual *i*, respectively. ε_i_ is the nuisance variable for individual *i*.

### Follicle count analysis

We sacrificed 15 sexually mature female mice, of which 10 were wildtype and 5 were knockout genotypes. Bilateral ovaries were fixed, sectioned to 5 μm, and stained with H&E. We performed morphological classification of follicles in both ovaries as described previously (Myers et al., 2004). We identified primordial follicles, primary follicles, secondary follicles, early antral follicles, antral follicles, preovulatory follicles, atretic follicles, and *corpora lutea.* In order to estimate the effect of genotype on gonad weight, we fit the following linear model:

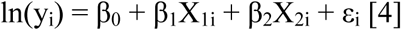

Where y_i_ is the number of total oocytes in bilateral ovaries of individual *i*, and X_1i_ and X_2i_ are genotype and age, respectively. ε_i_ is the nuisance variable for individual *i*.

### Breeding time analysis

We compiled comprehensive husbandry information over a period of greater than 1 year corresponding to 151 litters born representing all possible *Csmd1* wildtype, heterozygote, and knockout sire/dam breeding combinations. We calculated the number of days between first sire/dam co-habitation and birth of each litter. Next we subtracted an estimated C57BL/6 gestation time of 19 days (Murray et al., 2010) to estimate time to conception. We also calculated parental ages at conception. All density plots depicted in **Figure 4** reflect estimated time to conception for all 151 litters. In order to estimate the effect of maternal genotype on mating success, we controlled for paternal genotype by including wildtype sires only. We then fit the following linear model:

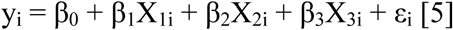

Where y_i_ is the estimated time to conception for mating pair *i*, X_1i_ is maternal genotype (wildtype, heterozygote, or knockout), X_2i_ is maternal age at conception, and X_3i_ is paternal age at conception. ε_i_ is the nuisance variable.

### Litter size analysis

We bred 44 females (8 wildtype, 27 heterozygote, and 9 homozygote) with 41 males (4 wildtype, 26 heterozygote, and 11 homozygote) over a period of 10 months to produce 99 litters, totaling 688 live births. All 9 parental genotype permutations [wt_dam_ x wt_sire_, wt_dam_ x het_sire_…hom_dam_ x hom_sire_] were represented multiple times (excepting het_dam_ x wt_sire_). We counted deaths in during the neonatal period (defined as 1-10 days by convention, although the vast majority of deaths occurred within 24-48 hours) and subtracted from the live birth total to obtain the final number of surviving pups (550 total). Next, we stratified each litter by maternal and paternal genotype status (*Csmd1* wildtype or heterozygous versus knockout) and fit the following linear model:

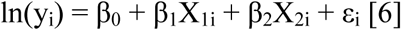

Where y_i_ is the number of surviving pups in litter i, and X_1i_ and X_2i_ are the maternal and paternal genotypes, respectively. ε_i_ is the nuisance variable for litter i.

### Mammary gland whole-mount analysis

Female littermates were collected at four developmental time points: (i) pre-pubescent (< 30 days of age); (ii) adult virgins; (iii) mid-pregnancy (estimated 14 days after copulation); (iv) post-weaning (7 days after weaning pups from mother’s nursing). Freshly-dissected whole inguinal mammary glands were fixed overnight in Carnoy’s solution (60% ethanol, 30% chloroform, 10% glacial acetic acid). Fixed tissues were washed and rehydrated in ethanol and water and stained in Carmine alum histological stain (0.5% Aluminium potassium sulphate, 0.2% Carmine) for 48 hours. Stained tissues were dehydrated with increasing concentrations of ethanol and stored in xylene to clear lipids for 48 hours. Finally, tissues were flattened mechanically and suspended in pure methyl salicylate prior to imaging. Due to the large size of whole mammary tissues, overlapping fields of view were captured and stitched together using the “Photomerge” function in Adobe Photoshop. Gaps in the backdrop of the merged images were filled using the “Content aware fill” function in Adobe Photoshop—if and only if the gaps did not overlap any portion of the tissue proper. All original images are available on the Conrad Lab website (http://genetics.wustl.edu/dclab/lee_et_al_images). To perform statistical comparison of duct morphology between genotypes, measurements of mammary gland ducts were derived from images using AngioTool64 v0.6a (Zudaire et al., 2011). First, a skeleton representation of the branched duct structure is generated from the input image, which is then used to compute a variety of morphological and spatial parameters for branching characterization. Since this software detects the branches by contrast on a black background, the images of whole mount mammary glands of adult mice were transformed into a compatible input using ImageJ 1.51n.

### Hormone measurements

We collected serum from 9 males (4 wildtype versus 5 knockout; mean age = 103 days) and 16 females in the proestrous stage (7 wildtype versus 9 knockout; mean age = 96 days) via submandibular collection. Each wildtype individual was matched with ≥ 1 knockout littermate. Female estrous cycle was determined by vaginal cytology, as described previously(Byers et al., 2012). All blood was drawn at approximately the same time of day, clotted for 90 minutes at room temperature, and centrifuged at 2000 x g for 15 minutes. Samples were stored at −20°C prior to hormone measurements. Male samples were quantified for LH/FSH (EMD Millipore) and testosterone (Immuno-Biological Laboratories Inc), and female samples were quantified for LH/FSH and estradiol (CALBIOTECH), as described by the University of Virginia Ligand Assay and Analysis Core (http://www.medicine.virginia.edu/research/institutes-and-programs/crr/lab-facilities).

### C3 deposition assay

Testes obtained post-dissection from *Csmd1* knockout and wild-type mice were decapsulated and washed in 1xPBS before mincing. Minced tissue was subjected to enzymatic dissociation as described by us previously (Lima et al, 2016). The crude cell preparation thus obtained was treated with ACK buffer (Life Technologies) for 5 min at room temperature to lyse erythrocytes present if any in the cell preparation. The isolated cells were incubated in αC3 (B9) primary antibody (Santa Cruz Biotechnology) for 45 minutes at room temperature (RT) diluted 1: 100 in FACS buffer (1x PBS, 5%FBS, 0.1% Sodium azide) along with 10% Fc block (to minimize non-specific binding and background fluorescence) followed by fluorophore tagged secondary antibody (1:250) incubation of 90 mins at RT in the dark with 3 washes of ice cold FACS buffer after each antibody incubation. Flow cytometry was performed with an Accuri C6 cytometer (BD Biosciences).

### *De novo* mutation calling

Whole genome sequencing and *de novo* mutation calling are described previously (Goldmann et al., 2016). Whole genome sequencing data were generated using the Complete Genomics Platform. All but one individual was excluded from each identical twin set in order to avoid double-counting same set of de novo mutations. After variant calling and QC we identified 2,058 DNMs across 709 trios. Finally, in order to assess the discrepancy, if any, between the high frequency of observed mutations about *CSMD1* and the intrinsic mutability of its primary sequence, we calculated a per-nucleotide mutation rate to every base across chromosome 8, based on pre-computed scores for 1,536 five-bp motifs.

### Sample-size estimation

Human genetic studies were carried out using existing datasets, 2/3 of which were generated by large epidemiological studies; thus, we simply used the sample sizes of cases and controls that were available to us. Based on empirical findings for diseases with similar genetic architecture (e.g. autism and schizophrenia) we hypothesized that sample sizes of approximately 500-1000 cases and thousands of controls would be sufficient to detect rare, large effect variants such as the 16p11 deletion that has a frequency of ~1% in cases of autism, which was originally detected as associated with only 180-500 cases of autism (Kumar et al., 2008; Weiss et al., 2008). For animal studies, we generated a large colony of wildtype, single and double knockouts that provided all phenotype and tissue data required for analyses. For tissue studies, we assayed at least 6 sections of the tissue of interest from at least 3 independent animals. For quantitative phenotyping, we assayed at least 3 independent animals (biological replicates). For ELISA (hormone assays) and FACS (protein abundance) experiments, a minimum of 3 technical replicates were taken on each animal and averaged.

## Author Contributions

D.F.C. devised the study. A.S.L. and D.F.C. led the experimental design. A.S.L. and D.F.C led the data analysis. A.S.L., A.C.L., N.H., K.A.V., W.S.W.W., R.E.W., J.P.A., and D.F.C. performed data analyses. J.E.N. supervised the data collection and sequencing of the human family trios. A.S.L., J.R., A.U., X.W., and R.A.H. performed experiments. A.S.L. and D.F.C. wrote the manuscript. All authors read and approved the manuscript.

## Description of Supplementary Data

Supplementary Figures-contains 6 supplemental figures and supplemental table.

In addition, there are two large tables of data that are provided separately:

Table S1 – The full set of CNV calls and inferred POI case/control status that we generated from the WHI SHARe cohort.

Table S2 – All deletions detected in introns 1-3 from SHARe, the azoospermia cohort, and UK Biobank, along with case/control status of each deletion carrier.

Table S3 – Results of breeding CSMD1 -/- C3 -/- double knockouts. Nineteen double knockouts were bred for 3-7 months. This table contains summary details on the outcome of breeding for each animal, including number of litters born and litter size(s).

## Acknowledgments

We thank all patients and study participants. D.F.C is supported by the National Institutes of Health (R01HD078641 and R01MH101810). A.S.L. is supported by a Distinguished Scholar Award from Washington University School of Medicine. The ITMI Premature Birth Study was funded by the Inova Health System. We thank Heather Lawson for training and consultation on animal husbandry. We thank Brianne Tabers for technical assistance with animal husbandry. We thank J. Carlson and S. Zollner for early access to Mr. Eel, software for annotating context-dependent mutation rates. We thank Kelle Moley and members of her laboratory (Praba Esakky, Michaela Reid, and Jessica Saben) for consultations on tissue preparations. We thank Tim Schedl and Nicole Rockweiler for helpful discussions. We thank Nicholas Ho for technical assistance with image quantification. We thank Jakob Goldmann and Christian Gilissen for providing us the software for calling and phasing the de novo mutations in family trio data. We thank Seungeun Lee for discussion of SKAT results. We thank Bill Eades and Chris Holley at the Washington University Siteman Flow Cytometry Core (NCI P30 CA91842) FACS services. We thank the Washington University Rheumatic Disease Core (NIHP30AR48335) for providing backcrossed mouse genotypes. We thank the University of Virginia Ligand Assay and Analysis Core (U54 DH28934) for hormone measurements.

## References

Baker, T.G. (1963). A Quantitative and Cytological Study of Germ Cells in Human Ovaries. Proc R Soc Lond B Biol Sci 158, 417–433.

Byers, S.L., Wiles, M.V., Dunn, S.L., and Taft, R.A. (2012). Mouse estrous cycle identification tool and images. PLoS One 7, e35538.

Chen, C.T., Fernandez-Rhodes, L., Brzyski, R.G., Carlson, C.S., Chen, Z., Heiss, G., North, K.E., Woods, N.F., Rajkovic, A., Kooperberg, C., and Franceschini, N. (2012). Replication of loci influencing ages at menarche and menopause in Hispanic women: the Women's Health Initiative SHARe Study. Hum Mol Genet 21, 1419-1432.

Circolo, A., Garnier, G., Fukuda, W., Wang, X., Hidvegi, T., Szalai, A.J., Briles, D.E., Volanakis, J.E., Wetsel, R.A., and Colten, H.R. (1999). Genetic disruption of the murine complement C3 promoter region generates deficient mice with extrahepatic expression of C3 mRNA. Immunopharmacology 42, 135-149.

Clarkson, R.W., Wayland, M.T., Lee, J., Freeman, T., and Watson, C.J. (2004). Gene expression profiling of mammary gland development reveals putative roles for death receptors and immune mediators in post-lactational regression. Breast Cancer Res 6, R92-109.

Day, F.R., Ruth, K.S., Thompson, D.J., Lunetta, K.L., Pervjakova, N., Chasman, D.I., Stolk, L., Finucane, H.K., Sulem, P., Bulik-Sullivan, B., Esko, T., Johnson, A.D., Elks, C.E., Franceschini, N., He, C., Altmaier, E., Brody, J.A., Franke, L.L., Huffman, J.E., Keller, M.F., et al. (2015). Large-scale genomic analyses link reproductive aging to hypothalamic signaling, breast cancer susceptibility and BRCA1-mediated DNA repair. Nat Genet 47, 1294-1303.

Day, F.R., Thompson, D.J., Helgason, H., Chasman, D.I., Finucane, H., Sulem, P., Ruth, K.S., Whalen, S., Sarkar, A.K., Albrecht, E., Altmaier, E., Amini, M., Barbieri, C.M., Boutin, T., Campbell, A., Demerath, E., Giri, A., He, C., Hottenga, J.J., Karlsson, R., et al. (2017). Genomic analyses identify hundreds of variants associated with age at menarche and support a role for puberty timing in cancer risk. Nat Genet 49, 834-841.

DeFalco, T., Potter, S.J., Williams, A.V., Waller, B., Kan, M.J., and Capel, B. (2015). Macrophages Contribute to the Spermatogonial Niche in the Adult Testis. Cell Rep 12, 1107-1119.

Escudero-Esparza, A., Kalchishkova, N., Kurbasic, E., Jiang, W.G., and Blom, A.M. (2013). The novel complement inhibitor human CUB and Sushi multiple domains 1 (CSMD1) protein promotes factor Imediated degradation of C4b and C3b and inhibits the membrane attack complex assembly. FASEB J 27, 5083-5093.

Foldi, C.J., Eyles, D.W., McGrath, J.J., and Burne, T.H. (2011). The effects of breeding protocol in C57BL/6J mice on adult offspring behaviour. PLoS One 6, e18152.

Friddle, C.J., Abuin, A., Ramirez-Solis, R., Richter, L.J., Buxton, E.C., Edwards, J., Finch, R.A., Gupta, A., Hansen, G., Holt, K.H., Hu, Y., Huang, W., Jaing, C., Key, B.W., Jr., Kipp, P., Kohlhauff, B., Ma, Z.Q., Markesich, D., Newhouse, M., Perry, T., et al. (2003). High-throughput mouse knockouts provide a functional analysis of the genome. Cold Spring Harb Symp Quant Biol 68, 311-315.

Gaytan, F., Morales, C., Bellido, C., Aguilar, E., and Sanchez-Criado, J.E. (1998). Ovarian follicle macrophages: is follicular atresia in the immature rat a macrophage-mediated event? Biol Reprod 58, 52-59.

Goldmann, J.M., Wong, W.S., Pinelli, M., Farrah, T., Bodian, D., Stittrich, A.B., Glusman, G., Vissers, L.E., Hoischen, A., Roach, J.C., Vockley, J.G., Veltman, J.A., Solomon, B.D., Gilissen, C., and Niederhuber, J.E. (2016). Parent-of-origin-specific signatures of de novo mutations. Nat Genet 48, 935-939.

Hays, J., Hunt, J.R., Hubbell, F.A., Anderson, G.L., Limacher, M., Allen, C., and Rossouw, J.E. (2003). The Women's Health Initiative recruitment methods and results. Ann Epidemiol 13, S18-77.

Hess, R.A., and Renato de Franca, L. (2008). Spermatogenesis and cycle of the seminiferous epithelium. Adv Exp Med Biol 636, 1-15.

Hoh Kam, J., Lenassi, E., Malik, T.H., Pickering, M.C., and Jeffery, G. (2013). Complement component C3 plays a critical role in protecting the aging retina in a murine model of age-related macular degeneration. Am J Pathol 183, 480-492.

Hotaling, J., and Carrell, D.T. (2014). Clinical genetic testing for male factor infertility: current applications and future directions. Andrology 2, 339-350.

Hsueh, A.J., Billig, H., and Tsafriri, A. (1994). Ovarian follicle atresia: a hormonally controlled apoptotic process. Endocr Rev 15, 707-724.

Huang, N., Wen, Y., Guo, X., Li, Z., Dai, J., Ni, B., Yu, J., Lin, Y., Zhou, W., Yao, B., Jiang, Y., Sha, J., Conrad, D.F., and Hu, Z. (2015). A Screen for Genomic Disorders of Infertility Identifies MAST2 Duplications Associated with Non-Obstructive Azoospermia in Humans. Biol Reprod.

Ingman, W.V., Wyckoff, J., Gouon-Evans, V., Condeelis, J., and Pollard, J.W. (2006). Macrophages promote collagen fibrillogenesis around terminal end buds of the developing mammary gland. Dev Dyn 235, 3222-3229.

Kamal, M., Shaaban, A.M., Zhang, L., Walker, C., Gray, S., Thakker, N., Toomes, C., Speirs, V., and Bell, S.M. (2010). Loss of CSMD1 expression is associated with high tumour grade and poor survival in invasive ductal breast carcinoma. Breast Cancer Res Treat 121, 555-563.

Kato, S., Shiratsuchi, A., Nagaosa, K., and Nakanishi, Y. (2005). Phosphatidylserine-and integrin-mediated phagocytosis of apoptotic luteal cells by macrophages of the rat. Dev Growth Differ 47, 153-161.

Kim, D., Pertea, G., Trapnell, C., Pimentel, H., Kelley, R., and Salzberg, S.L. (2013). TopHat2: accurate alignment of transcriptomes in the presence of insertions, deletions and gene fusions. Genome Biol 14, R36.

Kircher, M., Witten, D.M., Jain, P., O'Roak, B.J., Cooper, G.M., and Shendure, J. (2014). A general framework for estimating the relative pathogenicity of human genetic variants. Nat Genet 46, 310-315.

Korn, J.M., Kuruvilla, F.G., McCarroll, S.A., Wysoker, A., Nemesh, J., Cawley, S., Hubbell, E., Veitch, J., Collins, P.J., Darvishi, K., Lee, C., Nizzari, M.M., Gabriel, S.B., Purcell, S., Daly, M.J., and Altshuler, D. (2008). Integrated genotype calling and association analysis of SNPs, common copy number polymorphisms and rare CNVs. Nat Genet 40, 1253-1260.

Kraus, D.M., Elliott, G.S., Chute, H., Horan, T., Pfenninger, K.H., Sanford, S.D., Foster, S., Scully, S., Welcher, A.A., and Holers, V.M. (2006). CSMD1 is a novel multiple domain complement-regulatory protein highly expressed in the central nervous system and epithelial tissues. J Immunol 176, 4419-4430.

Kumar, R.A., KaraMohamed, S., Sudi, J., Conrad, D.F., Brune, C., Badner, J.A., Gilliam, T.C., Nowak, N.J., Cook, E.H., Jr., Dobyns, W.B., and Christian, S.L. (2008). Recurrent 16p11.2 microdeletions in autism. Hum Mol Genet 17, 628-638.

Kurilo, L.F. (1981). Oogenesis in antenatal development in man. Hum Genet 57, 86-92.

Laufer, J., Oren, R., Goldberg, I., Afek, A., Kopolovic, J., and Passwell, J.H. (1999). Local complement genes expression in the mammary gland: effect of gestation and inflammation. Pediatr Res 46, 608-612.

Lee, S., Emond, M.J., Bamshad, M.J., Barnes, K.C., Rieder, M.J., Nickerson, D.A., Team, N.G.E.S.P.-E.L.P., Christiani, D.C., Wurfel, M.M., and Lin, X. (2012). Optimal unified approach for rare-variant association testing with application to small-sample case-control whole-exome sequencing studies. Am J Hum Genet 91, 224-237.

Lesher, A.M., Zhou, L., Kimura, Y., Sato, S., Gullipalli, D., Herbert, A.P., Barlow, P.N., Eberhardt, H.U., Skerka, C., Zipfel, P.F., Hamano, T., Miwa, T., Tung, K.S., and Song, W.C. (2013). Combination of factor H mutation and properdin deficiency causes severe C3 glomerulonephritis. J Am Soc Nephrol 24, 53-65.

Li, J., Witten, D.M., Johnstone, I.M., and Tibshirani, R. (2012). Normalization, testing, and false discovery rate estimation for RNA-sequencing data. Biostatistics 13, 523-538.

Li, R., and Albertini, D.F. (2013). The road to maturation: somatic cell interaction and self-organization of the mammalian oocyte. Nat Rev Mol Cell Biol 14, 141-152.

Li, S., Zhou, W., Doglio, L., and Goldberg, E. (1998). Transgenic mice demonstrate a testis-specific promoter for lactate dehydrogenase, LDHC. J Biol Chem 273, 31191-31194.

Lie, P.P., Mruk, D.D., Lee, W.M., and Cheng, C.Y. (2010). Cytoskeletal dynamics and spermatogenesis. Philos Trans R Soc Lond B Biol Sci 365, 1581-1592.

Lima, A.C., Jung, M., Rusch, J., Usmani, A., Lopes, A., and Conrad, D.F. (2016). Multispecies Purification of Testicular Germ Cells. Biol Reprod 95, 85.

Lipkin, S.M., Moens, P.B., Wang, V., Lenzi, M., Shanmugarajah, D., Gilgeous, A., Thomas, J., Cheng, J., Touchman, J.W., Green, E.D., Schwartzberg, P., Collins, F.S., and Cohen, P.E. (2002). Meiotic arrest and aneuploidy in MLH3-deficient mice. Nat Genet 31, 385-390.

Liszewski, M.K., Farries, T.C., Lublin, D.M., Rooney, I.A., and Atkinson, J.P. (1996). Control of the complement system. Adv Immunol 61, 201-283.

Lopes, A.M., Aston, K.I., Thompson, E., Carvalho, F., Goncalves, J., Huang, N., Matthiesen, R., Noordam, M.J., Quintela, I., Ramu, A., Seabra, C., Wilfert, A.B., Dai, J., Downie, J.M., Fernandes, S., Guo, X., Sha, J., Amorim, A., Barros, A., Carracedo, A., et al. (2013). Human spermatogenic failure purges deleterious mutation load from the autosomes and both sex chromosomes, including the gene DMRT1. PLoS Genet 9, e1003349.

Luborsky, J.L., Meyer, P., Sowers, M.F., Gold, E.B., and Santoro, N. (2003). Premature menopause in a multi-ethnic population study of the menopause transition. Hum Reprod 18, 199-206.

Matzuk, M.M., and Lamb, D.J. (2008). The biology of infertility: research advances and clinical challenges. Nat Med 14, 1197-1213.

Murray, S.A., Morgan, J.L., Kane, C., Sharma, Y., Heffner, C.S., Lake, J., and Donahue, L.R. (2010). Mouse gestation length is genetically determined. PLoS One 5, e12418.

Myers, M., Britt, K.L., Wreford, N.G., Ebling, F.J., and Kerr, J.B. (2004). Methods for quantifying follicular numbers within the mouse ovary. Reproduction 127, 569-580.

Nelson, L.M. (2009). Clinical practice. Primary ovarian insufficiency. N Engl J Med 360, 606-614.

Ni, B., Lin, Y., Sun, L., Zhu, M., Li, Z., Wang, H., Yu, J., Guo, X., Zuo, X., Dong, J., Xia, Y., Wen, Y., Wu, H., Li, H., Zhu, Y., Ping, P., Chen, X., Dai, J., Jiang, Y., Xu, P., et al. (2015). Low-frequency germline variants across 6p22.2-6p21.33 are associated with non-obstructive azoospermia in Han Chinese men. Hum Mol Genet 24, 5628-5636.

Nusbaum, C., Mikkelsen, T.S., Zody, M.C., Asakawa, S., Taudien, S., Garber, M., Kodira, C.D., Schueler, M.G., Shimizu, A., Whittaker, C.A., Chang, J.L., Cuomo, C.A., Dewar, K., FitzGerald, M.G., Yang, X., Allen, N.R., Anderson, S., Asakawa, T., Blechschmidt, K., Bloom, T., et al. (2006). DNA sequence and analysis of human chromosome 8. Nature 439, 331-335.

O'Flynn O'Brien, K.L., Varghese, A.C., and Agarwal, A. (2010). The genetic causes of male factor infertility: a review. Fertil Steril 93, 1-12.

Ogundele, M.O. (1999). Anti-complement activities of human breast-milk. Inflamm Res 48, 437-445.

Perricone, R., Pasetto, N., De Carolis, C., Vaquero, E., Piccione, E., Baschieri, L., and Fontana, L. (1992). Functionally active complement is present in human ovarian follicular fluid and can be activated by seminal plasma. Clin Exp Immunol 89, 154-157.

Perry, J.R., Day, F., Elks, C.E., Sulem, P., Thompson, D.J., Ferreira, T., He, C., Chasman, D.I., Esko, T., Thorleifsson, G., Albrecht, E., Ang, W.Q., Corre, T., Cousminer, D.L., Feenstra, B., Franceschini, N., Ganna, A., Johnson, A.D., Kjellqvist, S., Lunetta, K.L., et al. (2014). Parent-of-origin-specific allelic associations among 106 genomic loci for age at menarche. Nature 514, 92-97.

Pollard, J.W. (2009). Trophic macrophages in development and disease. Nat Rev Immunol 9, 259-270.

Pompon, J., and Levashina, E.A. (2015). A New Role of the Mosquito Complement-like Cascade in Male Fertility in Anopheles gambiae. PLoS Biol 13, e1002255.

Riley-Vargas, R.C., Lanzendorf, S., and Atkinson, J.P. (2005). Targeted and restricted complement activation on acrosome-reacted spermatozoa. J Clin Invest 115, 1241-1249.

Schafer, D.P., Lehrman, E.K., Kautzman, A.G., Koyama, R., Mardinly, A.R., Yamasaki, R., Ransohoff, R.M., Greenberg, M.E., Barres, B.A., and Stevens, B. (2012). Microglia sculpt postnatal neural circuits in an activity and complement-dependent manner. Neuron 74, 691-705.

Schizophrenia Psychiatric Genome-Wide Association Study, C. (2011). Genome-wide association study identifies five new schizophrenia loci. Nat Genet 43, 969-976.

Schizophrenia Working Group of the Psychiatric Genomics, C. (2014). Biological insights from 108 schizophrenia-associated genetic loci. Nature 511, 421-427.

Sekar, A., Bialas, A.R., de Rivera, H., Davis, A., Hammond, T.R., Kamitaki, N., Tooley, K., Presumey, J., Baum, M., Van Doren, V., Genovese, G., Rose, S.A., Handsaker, R.E., Schizophrenia Working Group of the Psychiatric Genomics, C., Daly, M.J., Carroll, M.C., Stevens, B., and McCarroll, S.A. (2016). Schizophrenia risk from complex variation of complement component 4. Nature 530, 177-183.

Soumillon, M., Necsulea, A., Weier, M., Brawand, D., Zhang, X., Gu, H., Barthes, P., Kokkinaki, M., Nef, S., Gnirke, A., Dym, M., de Massy, B., Mikkelsen, T.S., and Kaessmann, H. (2013). Cellular source and mechanisms of high transcriptome complexity in the mammalian testis. Cell Rep 3, 2179-2190.

Steen, V.M., Nepal, C., Ersland, K.M., Holdhus, R., Naevdal, M., Ratvik, S.M., Skrede, S., and Havik, B. (2013). Neuropsychological deficits in mice depleted of the schizophrenia susceptibility gene CSMD1. PLoS One 8, e79501.

Stegle, O., Parts, L., Piipari, M., Winn, J., and Durbin, R. (2012). Using probabilistic estimation of expression residuals (PEER) to obtain increased power and interpretability of gene expression analyses. Nat Protoc 7, 500-507.

Sternlicht, M.D. (2006). Key stages in mammary gland development: the cues that regulate ductal branching morphogenesis. Breast Cancer Res 8, 201.

Stolk, L., Perry, J.R., Chasman, D.I., He, C., Mangino, M., Sulem, P., Barbalic, M., Broer, L., Byrne, E.M., Ernst, F., Esko, T., Franceschini, N., Gudbjartsson, D.F., Hottenga, J.J., Kraft, P., McArdle, P.F., Porcu, E., Shin, S.Y., Smith, A.V., van Wingerden, S., et al. (2012). Meta-analyses identify 13 loci associated with age at menopause and highlight DNA repair and immune pathways. Nat Genet 44, 260-268.

Sudlow, C., Gallacher, J., Allen, N., Beral, V., Burton, P., Danesh, J., Downey, P., Elliott, P., Green, J., Landray, M., Liu, B., Matthews, P., Ong, G., Pell, J., Silman, A., Young, A., Sprosen, T., Peakman, T., and Collins, R. (2015). UK biobank: an open access resource for identifying the causes of a wide range of complex diseases of middle and old age. PLoS Med 12, e1001779.

Tingen, C.M., Kiesewetter, S.E., Jozefik, J., Thomas, C., Tagler, D., Shea, L., and Woodruff, T.K. (2011). A macrophage and theca cell-enriched stromal cell population influences growth and survival of immature murine follicles in vitro. Reproduction 141, 809-820.

Tryka, K.A., Hao, L., Sturcke, A., Jin, Y., Wang, Z.Y., Ziyabari, L., Lee, M., Popova, N., Sharopova, N., Kimura, M., and Feolo, M. (2014). NCBI's Database of Genotypes and Phenotypes: dbGaP. Nucleic Acids Res 42, D975-979.

Wallace, W.H., and Kelsey, T.W. (2010). Human ovarian reserve from conception to the menopause. PLoS One 5, e8772.

Wang, K., Li, M., Hadley, D., Liu, R., Glessner, J., Grant, S.F., Hakonarson, H., and Bucan, M. (2007). PennCNV: an integrated hidden Markov model designed for high-resolution copy number variation detection in whole-genome SNP genotyping data. Genome Res 17, 1665-1674.

Wei, P.C., Chang, A.N., Kao, J., Du, Z., Meyers, R.M., Alt, F.W., and Schwer, B. (2016). Long Neural Genes Harbor Recurrent DNA Break Clusters in Neural Stem/Progenitor Cells. Cell 164, 644-655.

Weiss, L.A., Shen, Y., Korn, J.M., Arking, D.E., Miller, D.T., Fossdal, R., Saemundsen, E., Stefansson, H., Ferreira, M.A., Green, T., Platt, O.S., Ruderfer, D.M., Walsh, C.A., Altshuler, D., Chakravarti, A., Tanzi, R.E., Stefansson, K., Santangelo, S.L., Gusella, J.F., Sklar, P., et al. (2008). Association between microdeletion and microduplication at 16p11.2 and autism. N Engl J Med 358, 667-675.

White, J.K., Gerdin, A.K., Karp, N.A., Ryder, E., Buljan, M., Bussell, J.N., Salisbury, J., Clare, S., Ingham, N.J., Podrini, C., Houghton, R., Estabel, J., Bottomley, J.R., Melvin, D.G., Sunter, D., Adams, N.C., Sanger Institute Mouse Genetics, P., Tannahill, D., Logan, D.W., Macarthur, D.G., et al. (2013). Genome-wide generation and systematic phenotyping of knockout mice reveals new roles for many genes. Cell 154, 452-464.

Willott, G.M. (1982). Frequency of azoospermia. Forensic Sci Int 20, 9-10.

Wilson, T.E., Arlt, M.F., Park, S.H., Rajendran, S., Paulsen, M., Ljungman, M., and Glover, T.W. (2015). Large transcription units unify copy number variants and common fragile sites arising under replication stress. Genome Res 25, 189-200.

Yatsenko, A.N., Georgiadis, A.P., Ropke, A., Berman, A.J., Jaffe, T., Olszewska, M., Westernstroer, B., Sanfilippo, J., Kurpisz, M., Rajkovic, A., Yatsenko, S.A., Kliesch, S., Schlatt, S., and Tuttelmann, F. (2015). X-linked TEX11 mutations, meiotic arrest, and azoospermia in infertile men. N Engl J Med 372, 2097-2107.

Zhao, H., Xu, J., Zhang, H., Sun, J., Sun, Y., Wang, Z., Liu, J., Ding, Q., Lu, S., Shi, R., You, L., Qin, Y., Zhao, X., Lin, X., Li, X., Feng, J., Wang, L., Trent, J.M., Xu, C., Gao, Y., et al. (2012). A genome-wide association study reveals that variants within the HLA region are associated with risk for nonobstructive azoospermia. Am J Hum Genet 90, 900-906.

Zudaire, E., Gambardella, L., Kurcz, C., and Vermeren, S. (2011). A computational tool for quantitative analysis of vascular networks. PLoS One 6, e27385.

